# Cell type-specific gene expression dynamics during human brain maturation

**DOI:** 10.1101/2023.09.29.560114

**Authors:** Christina Steyn, Ruvimbo Mishi, Stephanie Fillmore, Matthijs B. Verhoog, Jessica More, Ursula K. Rohlwink, Roger Melvill, James Butler, Johannes M. N. Enslin, Muazzam Jacobs, Tatjana Sauka-Spengler, Maria Greco, Sadi Quiñones, Chris G. Dulla, Joseph V. Raimondo, Anthony Figaji, Dorit Hockman

**Affiliations:** Division of Cell Biology, Department of Human Biology, University of Cape Town, Cape Town, South Africa; Neuroscience Institute, University of Cape Town, Cape Town, South Africa; Division of Neurosurgery, Department of Surgery, University of Cape Town, Cape Town, South Africa; Division of Neurology, Department of Medicine, University of Cape Town, Cape Town, South Africa; Institute of Infectious Disease and Molecular Medicine, University of Cape Town, Cape Town, South Africa; Division of Immunology, Department of Pathology University of Cape Town; National Health Laboratory Service, South Africa; Radcliffe Department of Medicine, MRC Weatherall Institute of Molecular Medicine, University of Oxford, Oxford, UK; Stowers Institute for Medical Research, Kansas City, MO, USA; Single Cell Facility, MRC Weatherall Institute of Molecular Medicine, University of Oxford, Oxford, UK; Department of Neuroscience, Graduate School of Biomedical Sciences, Tufts University School of Medicine, Boston, MA, USA; Graduate School of Biomedical Science, Tufts University School of Medicine, Boston, MA, USA

## Abstract

The human brain undergoes protracted post-natal maturation, guided by dynamic changes in gene expression. Most studies exploring these processes have used bulk tissue analyses, which mask cell type-specific gene expression dynamics. Here, using single nucleus (sn)RNA-seq on temporal lobe tissue, including samples of African ancestry, we build a joint paediatric and adult atlas of 75 cell subtypes, which we verify with spatial transcriptomics. We explore the differences between paediatric and adult cell types, revealing the genes and pathways that change during brain maturation. Our results highlight excitatory neuron subtypes, including the *LTK* and *FREM* subtypes, that show elevated expression of genes associated with cognition and synaptic plasticity in paediatric tissue. The new resources we present here improve our understanding of the brain during its development and contribute to global efforts to build an inclusive brain cell map.

## Introduction

The adult human brain is a complex assembly of diverse cell types, which has been defined with unprecedented accuracy using single cell transcriptomics^1–4^. This adult transcriptomic signature is set up over a protracted period of development, which begins in the embryo and continues after birth. While the single cell diversity of the embryonic human brain has been explored^5,6^, little is known about how these cell type-specific gene expression profiles change during childhood^7^. Most existing studies have used bulk transcriptomic approaches, which revealed a dramatic period of global gene expression change during the late foetal/early infancy transition, that stabilises during childhood (1 to <12-years-old) and adolescence (12 to <20-years-old)^6,8–11^. Bulk transcriptomics, however, cannot reveal the more subtle, cell type-specific changes in gene expression that drive brain maturation from childhood, through adolescence to adulthood.

Childhood and adolescence are periods of important changes in brain structure, during which neuronal connections are refined and strengthened. While synaptogenesis peaks in the early postnatal period, synaptic pruning activity begins during late childhood, peaks during adolescence, and then gradually decreases^12–14^. These stages therefore represent periods of enhanced susceptibility to environmental influence, as well as increased neuropsychiatric risk^15^. Describing the typical cell type-specific gene expression trajectories of the maturing brain will allow us to assess the effects of genetic perturbations and early adverse experiences on brain maturation. Furthermore, investigating the driving forces behind cell type-specific maturational processes may help develop targeted therapies for neurological disease^16^.

To this end, the Paediatric Cell Atlas (PCA)^17^, a branch of the Human Cell Atlas (HCA)^3^, aims to ensure that the benefits of single cell transcriptomics are available to children as well as adults from diverse populations^3,17^. Africa has the most genetically diverse^18^ and youngest population^19^ worldwide and by 2050, 37% of the world’s children will grow up in Africa^20^. Consequently, it is essential to include the African paediatric population in the PCA’s efforts. A reference paediatric brain cell atlas that includes data from African donors will contribute to developing treatments for locally prevalent conditions, such as tuberculosis meningitis (TBM) and HIV^21,22^. In addition, studying the differences in gene expression dynamics between adult and paediatric brains may explain why the manifestation of neurological conditions and responses to therapies differ across the lifespan^17^.

To contribute to these endeavours, we present a joint paediatric and adult temporal cortex cell atlas, including samples from eight Southern African donors, annotated using the Allen Brain Map middle temporal gyrus (MTG) cell taxonomy^1^. We validate our annotation using spatial transcriptomics analysis. In addition, we use *de novo* marker gene analysis with machine learning tools to compare our paediatric and adult datasets to the existing MTG cell taxonomy and compare markers that define paediatric versus adult cell states. Using differential gene expression analysis, we highlight 21 cell subtypes that show differential expression of genes involved in neurodevelopment and cognition. Finally, we use our datasets to define the cell type-specific gene expression of putative site-of-disease TBM biomarkers^23^. Overall, we highlight the subtle cell type-specific differences between the paediatric and adult brain and expand the representation of diverse paediatric populations in the HCA.

## Results

### A joint paediatric and adult temporal cortex cell atlas

We generated snRNA-seq libraries from five paediatric and three adult donor temporal cortex tissue samples. The majority of our samples were obtained from surgeries to treat epilepsy (Extended Data Table 1). These new libraries were analysed alongside similar published datasets^24^, resulting in a total of 23 snRNA-seq datasets (including technical replicates) from 12 individuals (six paediatric and six adult) (Fig. 1a). The samples were sequenced to a median depth of 19,853 reads per nucleus, with 176,012 nuclei remaining after removing low quality barcodes (Extended Data Fig. 1, Extended Data Table 2). While our new datasets had a lower average sequencing depth than the co-analysed published datasets, the average number of genes and transcripts detected across datasets was similar (Extended Data Table 2).

**Fig. 1:**
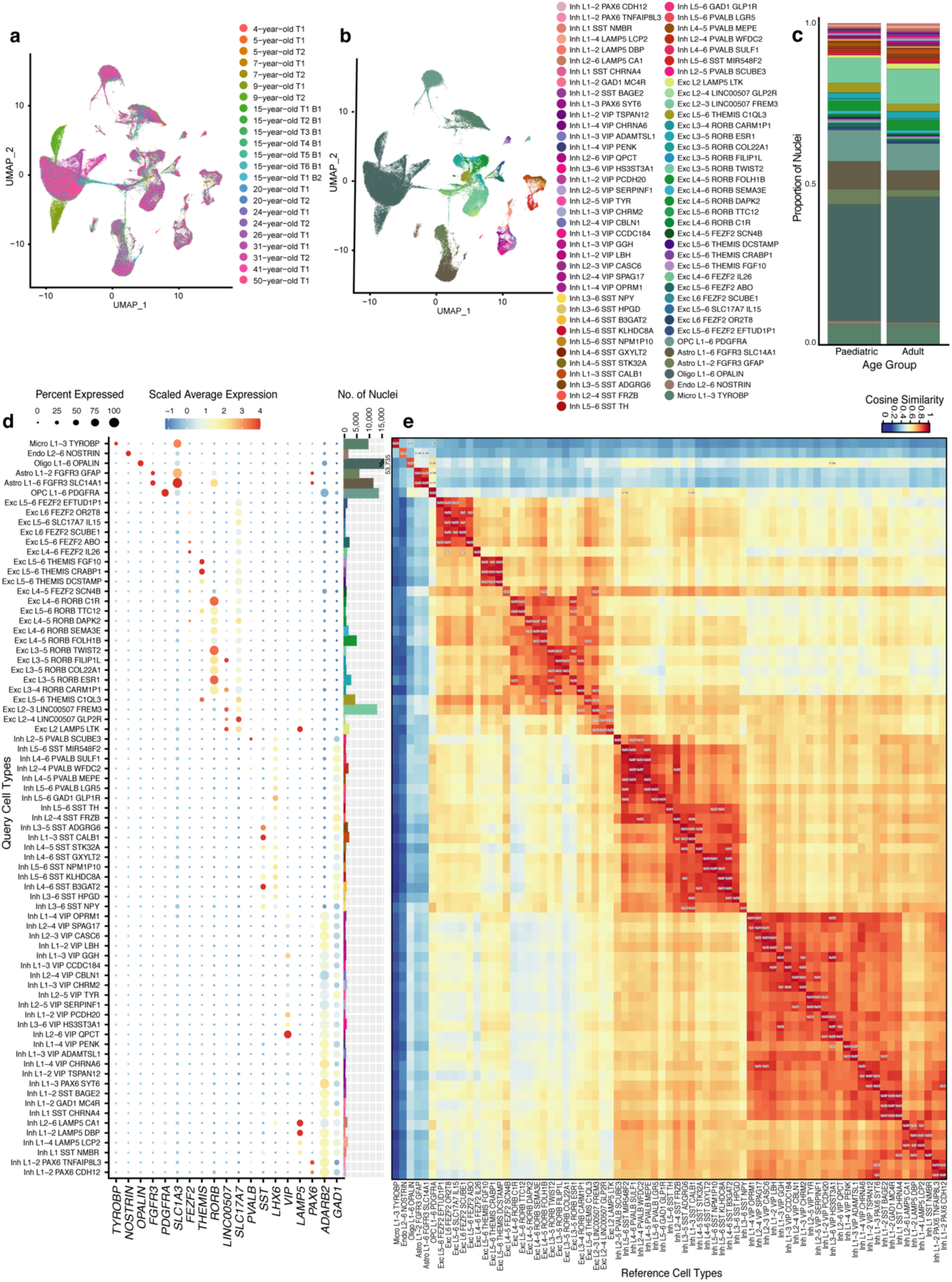
Annotation of nuclei by label transfer identifies 75 cell types across the 23 datasets. **a**, Data integration shows alignment of nuclei across the technical (T) and biological (B) replicates from donors ranging in age from 4 to 50 years. **b**, UMAP plot annotated to show the 75 cell types from the Allen Brain Map MTG atlas after filtering to retain nuclei with high confidence annotations. Each cell type is annotated with 1) a major cell class (e.g. Exc for excitatory neuron), 2) the cortical layer the cell is associated with (e.g. L2 for layer 2), 3) a subclass marker gene and 4) a cluster-specific marker gene. **c**, Stacked barplot showing the proportion of nuclei per cell type for each age category out of the total number of nuclei for each group. The cell types are coloured as in **b.** See Extended Data Table 3 for details of statistical tests performed. **d**, Validation of the high-resolution cell type annotations shows a high degree of correspondence in the expression of known cell type-specific marker genes (x axis) with their expected cell type (y axis) (left). The number of nuclei per cell type is shown on the right. **e**, Correlation plot showing the cosine similarity scores assessing similarity between the annotated cell types in our dataset (y axis as in **d**) and the MTG reference dataset (x axis) based on the log normalized expression counts of the top 2000 shared highly variable features between query and reference datasets.

Using data integration and clustering we aligned similar cell types across the 23 datasets, yielding 40 clusters (Fig. 1a, Extended Data Fig. 1i-h). Each cluster was assigned to one of the major brain cell types (level 1 annotation) based on marker gene expression (Extended Data Fig. 2a, Extended Data Table 3, Supplementary Figure 1). Additionally, we used label transfer^25^ to classify each nucleus according to the Allen Brain Map MTG atlas^1^ (level 2 annotation) (Extended Data Table 3). Barcodes with discordant level 1 and level 2 annotations (17.94%) were removed to focus downstream analyses on nuclei with high confidence annotations (Extended Data Table 3). Based on marker gene analysis^1^ (Extended Data Fig. 2b), many of these filtered barcodes are likely multiplets or nuclei contaminated with ambient mRNA.

All 75 reference cell types were present in the final filtered dataset of 144,438 nuclei (Fig. 1b; Extended Data Table 3) and expressed the expected marker genes^1^ (Fig. 1d). Both neuronal and non-neuronal cell types showed high correlation with the corresponding reference cell types^1^ (cosine similarity score > 0.83) and lower correlation to other subtypes within their class (Fig. 1e). This pattern was maintained when we considered either the paediatric or adult datasets on their own, with the majority of paediatric cell types showing only slightly lower similarity scores than the adults (Extended Data Table 3), which is likely due to the reference dataset only containing adult data. The cell composition of the samples was very similar with no significant differences in cell type proportions between paediatric and adult samples or between biological sexes (Fig. 1c; Extended Data Figure 2c-d; Extended Data Table 3). Similar to the reference atlas^1^, oligodendrocytes were the most common non-neuronal cell type and Exc_L2-3_LINC00507_FREM3 was the most common neuronal subtype. Neuronal clusters had a greater number of expressed genes and unique molecular identifiers (UMIs) compared to non-neuronal cells (Extended Data Figure 3a), while excitatory neurons had a greater number of genes detected per nucleus than inhibitory neurons (Extended Data Table 3). When comparing the paediatric to adult cell types, there were no significant differences in the number of genes or UMIs between the age categories. Overall, the quality and composition of the paediatric and adult cell atlases were very similar.

### Spatial mapping of cell types reveals similar tissue cytoarchitecture in adult and paediatric temporal cortex

Next, we used spatial transcriptomics to explore the positions of our annotated cell types within the temporal cortex. We generated Visium datasets from adult (31-year-old) and paediatric (15-year-old) temporal cortex samples (two sections each; Extended Data Table 1; Extended Data Fig. 4). The four Visium libraries were sequenced to a median depth of 87,178 reads per spot (median of 5,878 UMIs and 2,745 genes per spot) (Extended Data Table 4).

Using *cell2location*^26^, we calculated cell type abundance estimates for each Visium spot, with our annotated snRNA-seq dataset as a reference. Oligodendrocytes were the most common cell type, while Exc_L2_LAMP5_LTK was the most abundant neuronal cell type (Extended Data Fig. 5a). The annotated cell types mapped to their expected cortical layer locations across all tissue sections (Fig. 2a; Extended Data Fig. 5b), matching the spatial expression of known cortical layer marker genes^1,27,28^ (Fig. 2b). These layered expression patterns were verified for a subset of layer-specific marker genes using *in situ* hybridisation (Extended Data Fig. 6).

**Fig. 2:**
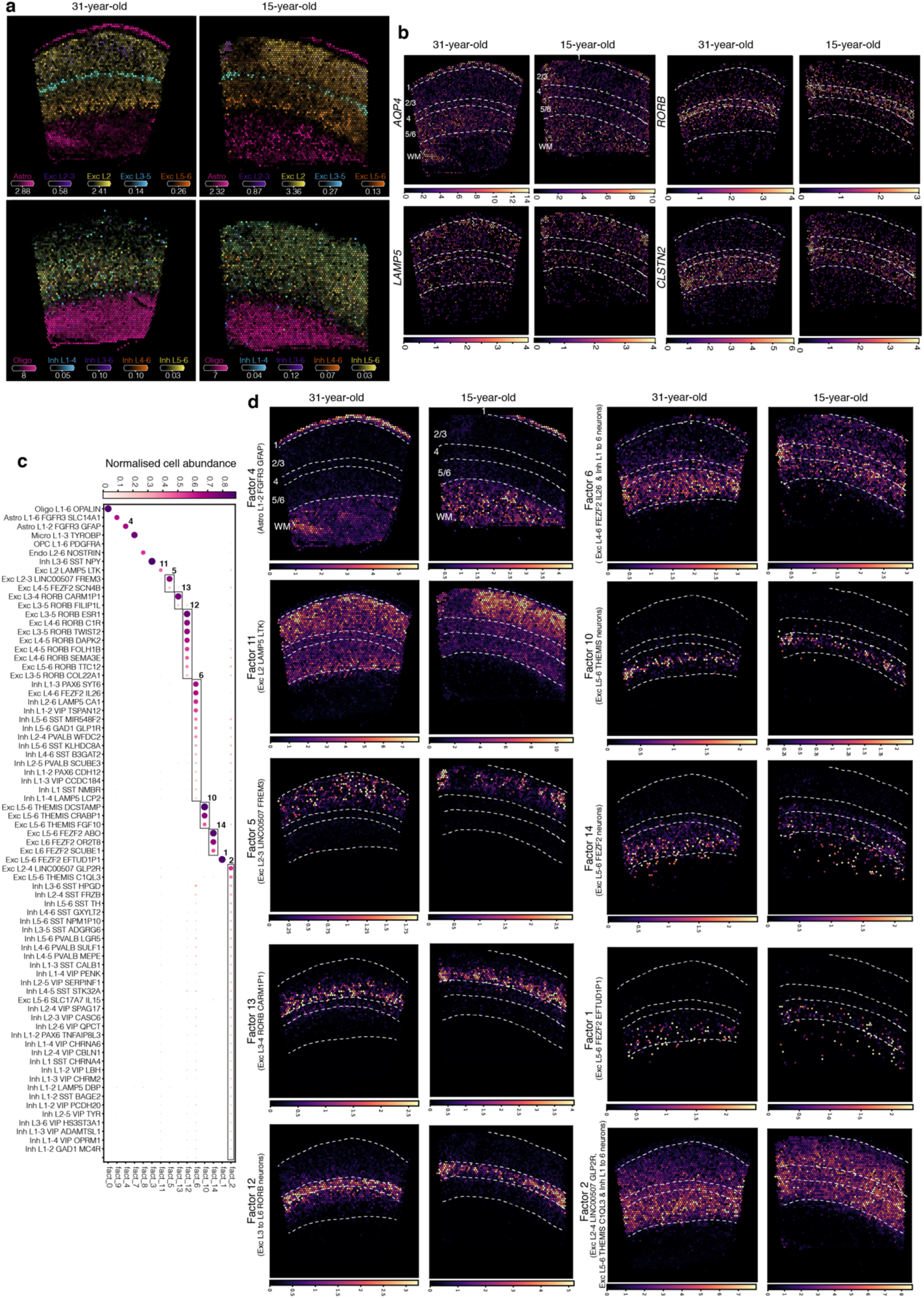
Visium spatial transcriptomics in the adult and paediatric temporal cortex validates snRNA-seq annotation. **a**, Estimated cell type abundances (colour intensity) in the 31-year-old and 15-year-old temporal cortex tissue sections for a selection of cell types including non-neuronal cell types, excitatory neurons (top row) and inhibitory neurons (bottom row). **b**, Visium gene expression profiles (colour intensity) for a selection of known cortical layer marker genes in the 31-year-old and 15-year-old temporal cortex tissue sections including *AQP4* (layer 1), *LAMP5* (layer 2), RORB (layer 4) and *CLSTN2* (layer 5-6). **c**,**d**, Identification of co-locating cell types using NMF. The dot plot (**c**) shows the NMF weights of the cell types (rows) across each of the NMF factors (columns), which correspond to tissue compartments. Block boxes indicate cell types that co-locate within the indicated compartments. Spatial plots show (**d**) show the NMF weights for selected NMF factor/tissue compartment across the 31-year-old and 15-year-old temporal cortex tissue sections. Panels are displayed in the same order as the dotplot in (**c**), with the dominant cell types for each factor indicated in brackets. Dashed white lines and numbers indicate estimated cortical layer boundaries as indicated in the first two panels of **b** and **d**. WM: white matter. See also Extended Data Figs 4-6.

To examine the co-location of cell types within the layered structure of the temporal cortex, non-negative matrix factorization (NMF) was performed resulting in 15 cellular compartments, which were visualised across the Visium samples, revealing their spatial distribution (Fig. 2c-d, Extended Data Fig. 5c). In both the paediatric and adult datasets, there was clear co-location of the expected neuronal cell types within overlapping compartments across the cortical layers. Layer 2 was dominated by Exc_L2_LAMP5_LTK (factor_11) and Exc_L2-3_LINC00507_FREM (factor_5), layer 3 by Exc_L3-4_RORB_CARM1P1 (factor_13), layer 4 by the *RORB* excitatory neuron subtypes (factor_12), layer 5 by the *THEMIS* excitatory neuron subtypes (factor_10) and layer 6 by the *FEZF2* excitatory neuron subtypes (factor_14 and factor_1), with the latter extending into the white matter. Inhibitory neurons were primarily associated with factors 6 and 2, which were more widely spread across the layers. Interestingly, these factors were more strongly associated with layers 5/6 in the adult than in the paediatric samples. The two astrocyte subtypes were confirmed to have distinct distribution profiles, with Astro_L1-2_FGFR3_GFAP (factor_4) located primarily in layer 1 and the white matter, and Astro_L1-6_FGFR3_SLC14A1 (factor_9) more widely distributed. The remaining non-neuronal cell types were largely associated with factors located in layer 1 and the white matter.

Overall, our spatial transcriptomic analyses provide support for our annotation approach, showing the expected spatial distribution of annotated cell types, and revealing a largely similar tissue cytoarchitecture in adult and paediatric temporal cortex tissue.

### A machine learning approach identifies new temporal cortex cell type markers

To establish a standardized approach for defining cell types, it has been proposed to use the minimum combination of gene markers that can classify a cell type and distinguish it from other cell types^29,30^. Towards achieving this, Aevermann et al. (2021)^29^ developed the machine learning tool, *NS-Forest V2.0*, which they applied to the MTG cell atlas. Ideally, these MTG minimal markers would be conserved in similar datasets to facilitate accurate comparisons across different studies^31^. Indeed, we found that the majority of MTG cell atlas minimal markers^29^ (∼94%) are expressed at significantly higher levels in the expected cell types than in other cell types (Extended Data Fig. 7, Extended Data Table 5).

Application of the *NS-Forest V2.0*^29^ algorithm to our down-sampled snRNA-seq datasets (see Methods) revealed 202 paediatric and 196 adult minimal marker genes (Fig. 3; Extended Data Table 5). The median F-beta score per cell type (the measure of the discriminative power of a given combination of marker genes; paediatric=0.55; adult=0.6) and the average binary expression score (a measure of an individual gene’s classification power; paediatric=0.9; adult=0.89) were comparable across age groups and only slightly lower than that obtained for the MTG cell atlas (0.68 and 0.94 respectively)^29^. 47 paediatric (23.3%) and 45 adult (23.0%) minimal markers overlapped with existing markers^29^ (Fig. 3; Extended Data Table 5). However, there was a greater overlap in minimal markers between the paediatric and adult datasets, with 68 markers (∼34%) present in both lists. MERFISH^32^ spatial transcriptomic analysis of a subset of minimal makers that were shared between paediatric and adult datasets confirmed their co-expression with previously described minimal markers^29^ in adult (31-year-old) and paediatric (15-year-old) temporal cortex samples (Extended Data Fig. 8).

**Fig. 3:**
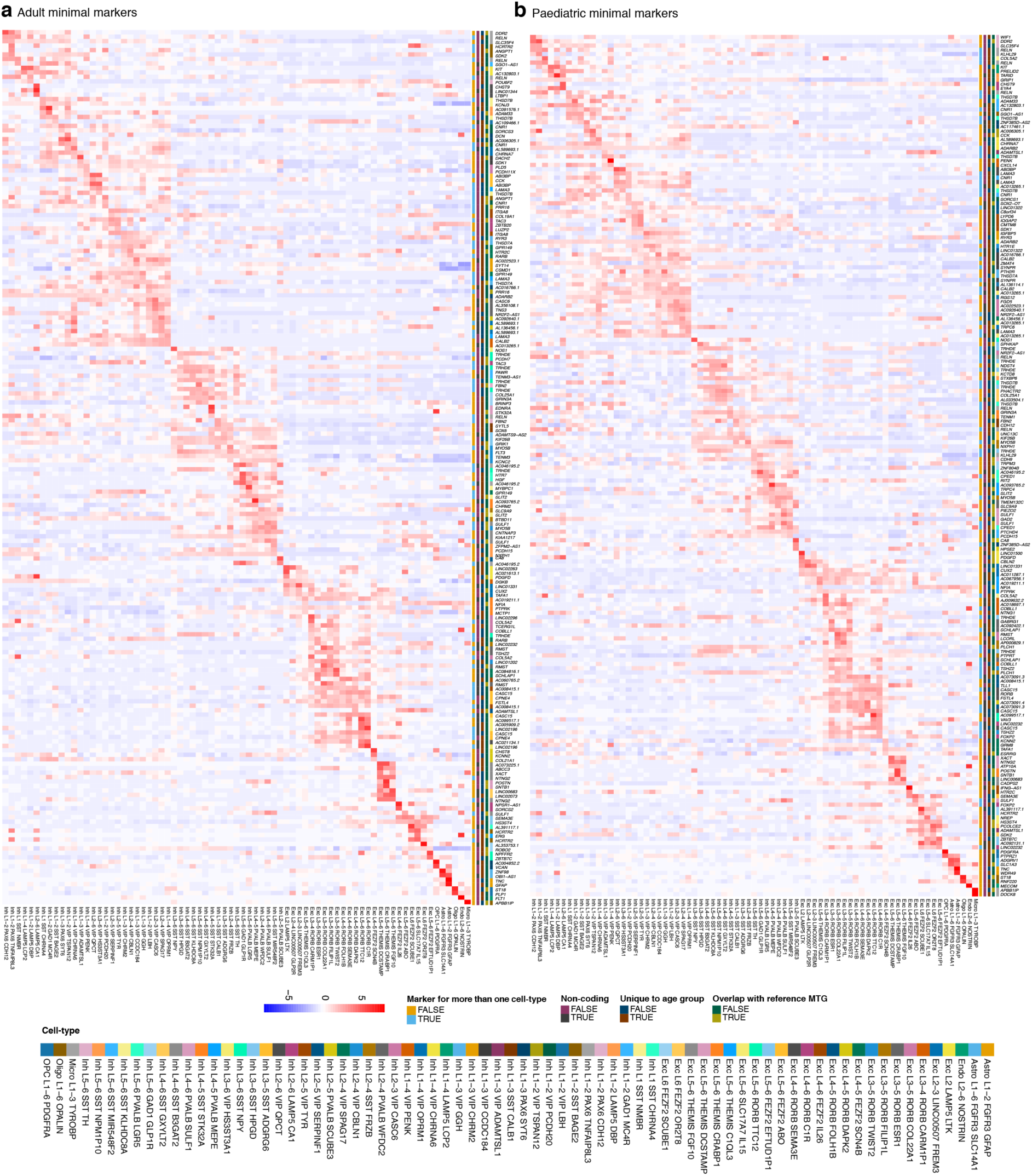
NS-Forest identifies minimal marker genes distinguishing the cell types in the paediatric and adult temporal cortex snRNA-seq datasets. **a**,**b**, Heatmap showing the scaled average normalised expression counts of the NS-Forest minimal marker genes (y-axis) identified for 75 cortical cell types (x-axis) across the six adult (**a**) and six paediatric (**b**) datasets. As input into NS-Forest, the nuclei of each sample were randomly down-sampled to the size of the sample with the fewest nuclei. Heatmaps show gene expression values for the down-sampled datasets. The minimal marker genes are annotated (colour codes on the y-axes) according to whether they are unique to a given cell type, whether they are coding/non-coding genes, whether they are unique to the indicated age group, whether they overlap with existing MTG minimal marker gene sets for the same cell type, and according to the cell type they define.

Our minimal marker analysis revealed improved markers for some cell types when compared to the reference MTG cell atlas. In our datasets, the long non-coding RNA, *LINC01331*, is a minimal marker for Exc_L2-3_LINC00507_FREM3 with a beta score of 1, indicating high specificity. In contrast, one of the existing markers for this cell type, *PALMD*, is more highly expressed in endothelial cells in our datasets (Fig. 3; Extended Data Fig. 9a-b). This discrepancy is likely due to the lower percentage of endothelial cells in the MTG cell atlas compared to our datasets (0.06% vs 0.9%)^1^. Similarly, one of the existing MTG cell atlas markers for Exc_L5-6 _THEMIS_CRABP1, *OLFML2B*, is more highly expressed in other layer 5/6 neurons in our dataset, whereas our minimal marker, *POSTN*, shows greater specificity (Fig. 3; Extended Data Fig. 9c-d). Additionally, UMAP analysis of our annotated datasets using our minimal marker gene list for each age group, in comparison to an equivalent number of random genes, resulted in better grouping of the cell subtypes into clusters, similar to the original UMAP plot (compare Fig. 1a and Fig. 4 a-b). This analysis reveals that our shortlists of ∼200 marker genes capture much of the underlying transcriptomic diversity in our datasets.

**Fig. 4:**
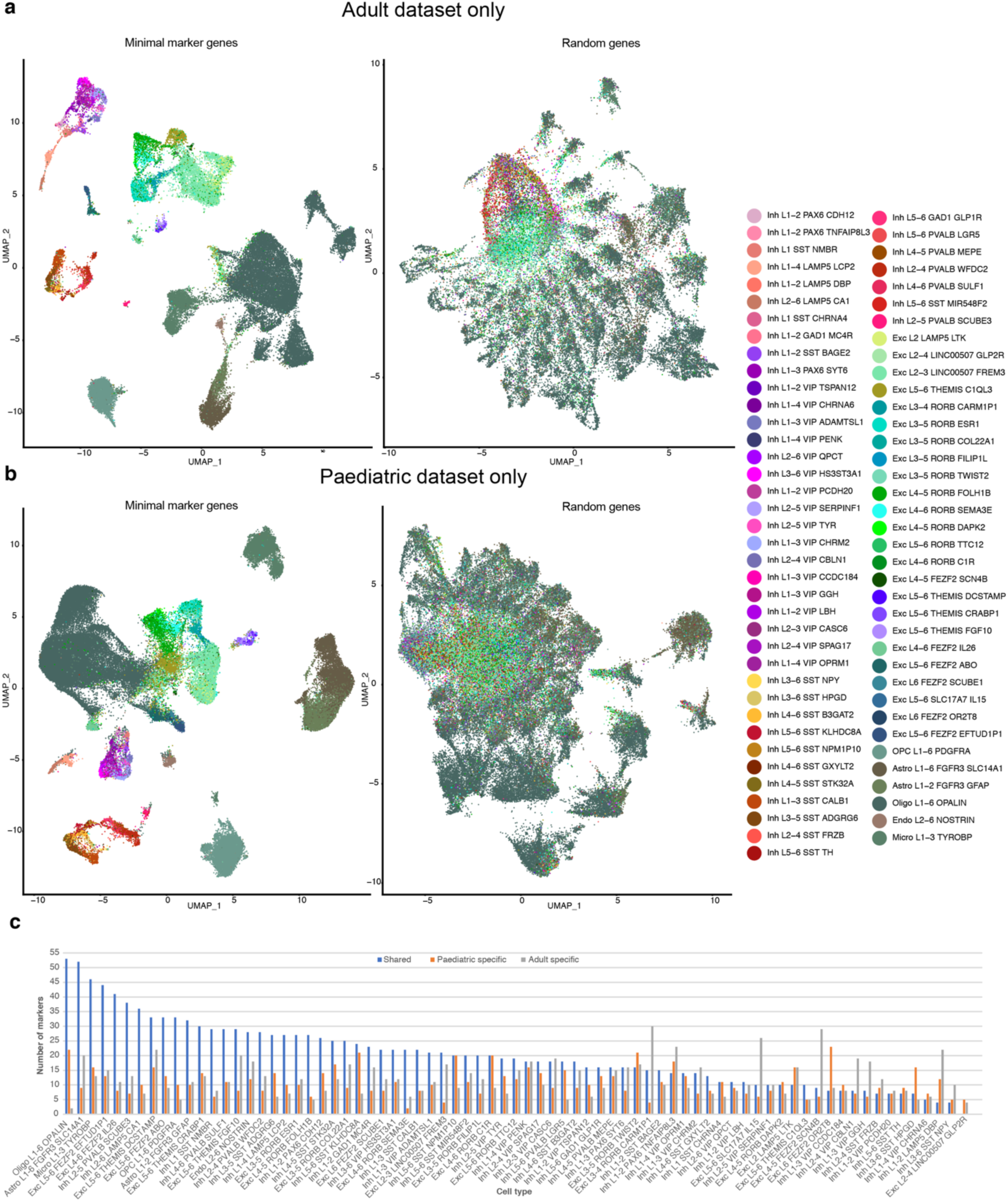
Validation of NS-Forest minimal markers and assessment of the top NS-forest markers. **a**,**b**, Annotated UMAP plots following data integration using either the minimal marker genes (left) or the equivalent number of a random set of genes (right) as anchors for the adult (**a**) and paediatric (**b**) datasets. The colour scheme for the cell types is in accordance with the MTG cell taxonomy. **c**, Overlap of the paediatric and adult NS-Forest markers with a high binary expression score (> 0.7) per cell type. The bar plot shows the number of shared markers between paediatric and adult datasets (blue), the number of markers unique to the paediatric datasets (orange), and the number of markers unique to the adult datasets (grey) for each cell type.

Gene ontology (GO) analysis of our minimal marker gene lists revealed significant enrichment of GO terms related to development, cell signalling, extracellular matrix and synapse organisation, when considering the paediatric and adult datasets individually or together (Extended Data Table 6). These results suggest that genes involved in neuronal development and signalling are key to neuronal identity as the brain matures and in adult life. To further assess the difference in cell type markers between our paediatric and adult datasets, we expanded our analysis to include all genes with a high NS-Forest binary expression score (> 0.7)^29^. For most cell types, the majority of these top markers (>18 genes) were shared between our paediatric and adult datasets (Fig. 4c; Extended Data Tables 7-8). The oligodendrocytes showed the highest number of shared marker genes (53), as well as the second highest number of paediatric-specific markers (22). Exc_L3-4_RORB_CARM1P1 had the highest number of adult-specific marker genes (30), while Exc_L2-4_LINC00507_GLP2R had no shared markers. GO analysis of the shared oligodendrocyte marker genes revealed driver terms related to oligodendrocyte structure and function, including “structural constituent of myelin sheath”, while the top driver terms for the paediatric-specific markers were “oligodendrocyte differentiation” and “myelination” (Extended Data Table 6). Overall, our expanded marker gene analysis suggests that neuronal cell types show greater dissimilarity between their paediatric and adult states than non-neuronal cells. It is likely that more diversity in the non-neuronal marker gene profiles could be revealed with subdivision into further subtypes.

### Differential gene expression analysis highlights enriched expression of genes associated with neurodevelopment in paediatric samples

To identify genes that were upregulated in the paediatric cell populations and thus might be involved in brain maturation, we conducted cell type-specific differential gene expression analysis with *DESeq2*^33^. In total, we detected 165 significantly differentially expressed genes (DEGs) across 21 cell types (123 upregulated in paediatric samples and 42 downregulated), with some DEGs associated with multiple cell types (Fig. 5a; Extended Data Table 9-10). For all DEGs, the change in expression was accompanied by a corresponding change in the percentage of nuclei expressing the gene (Extended Data Table 10). *BayesSpace*^34^ analysis of a subset of DEGs in our Visium datasets confirmed that the genes were expressed at higher levels in the 15-year-old compared to the 31-year-old (Extended Data Fig. 10).

**Fig. 5:**
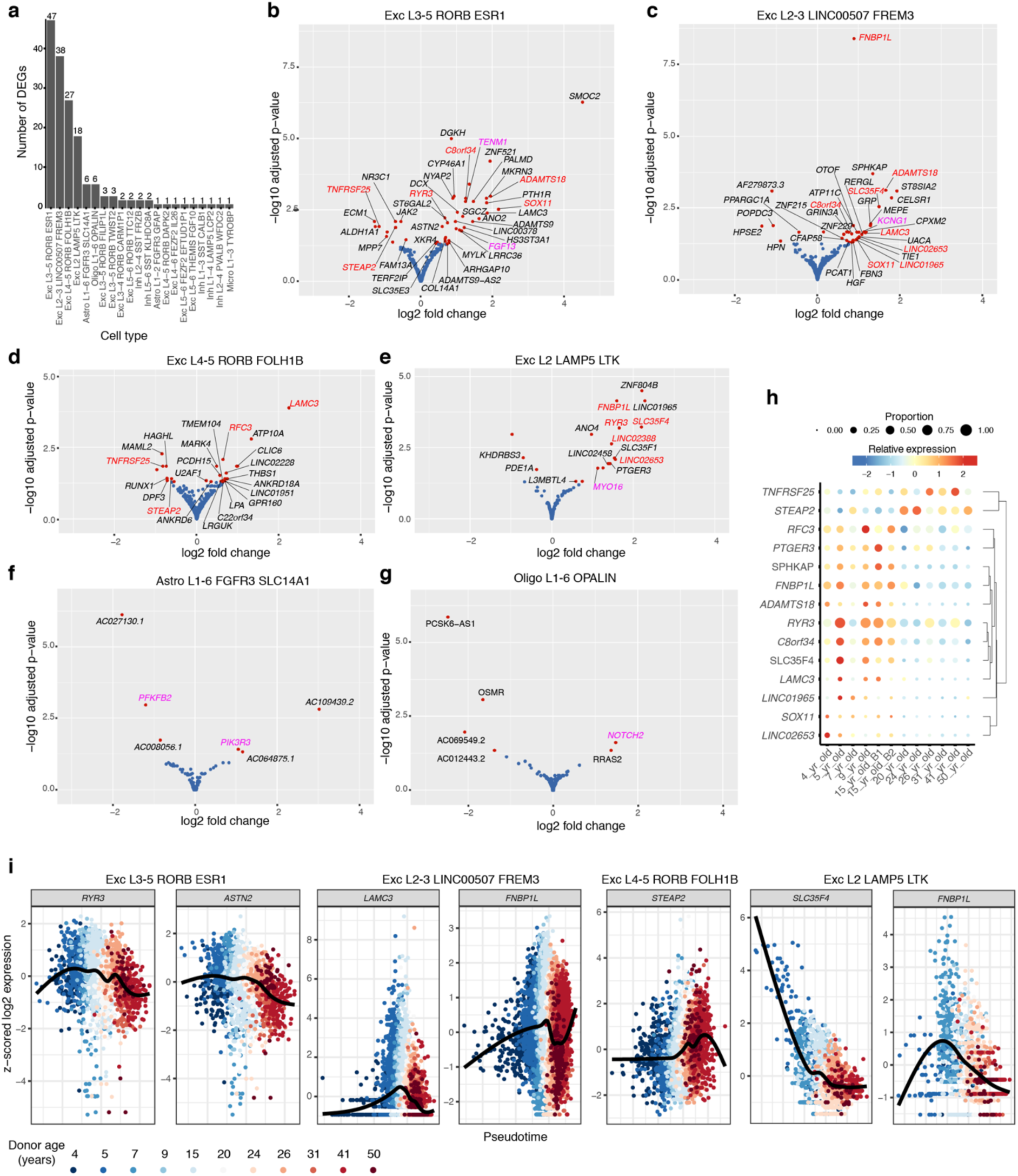
Differential expression analysis reveals genes guiding temporal cortex maturation. **a**, 21 cell types with DEGS. **b-g**, Volcano plots showing log_2_FoldChange (x axis) and -log_10_padj values (y axis) for all DESeq2-tested genes in Exc_L3-5_RORB_ESR1, Exc L2-3_LINC00507_FREM3, Exc_L4-5_RORB_FOLH1B, Exc_L2_LAMP5_LTK, Astro_L1-6_FGFR3_SLC14A1 and Oligo L1-6 OPALIN. Red dots indicate genes that were significantly upregulated or downregulated in paediatric samples (padj<0.05 & abs(log_2_FoldChange)>10%) and selected genes are labelled. Red labels indicate DEGs shared between neuronal cell types. Magenta labels indicate DEGS not shared between cell types that are discussed in the text. Blue dots indicate non-significant genes (padj>0.05 or abs(log_2_FoldChange)<10%). **h**, Dot plot showing the scaled average normalised expression across samples for DEGS shared between Exc_L3-5_RORB_ESR1, Exc L2-3_LINC00507_FREM3, Exc_L4-5_RORB_FOLH1B, Exc_L2_LAMP5_LTK, Exc_L3-4_RORB_CARM1P1 and Exc_L3-5_RORB_FILIP1L**. i**, psupertime gene expression trajectories for selected DEGs in the indicated cell types. The *x*-axis is the calculated psupertime value for each cell, coloured by sample of origin. The black lines are smoothened curves fit by geom_smooth in the R package ggplot2.

Many of the excitatory neuron subtypes shared DEGs that are known to be developmentally regulated in the mammalian brain (Fig. 5b-e,h). *LAMC3*, a subunit of the extracellular matrix protein laminin, was upregulated in three paediatric subtypes (Exc_L3-5_RORB_ESR1, Exc_L2-3_LINC00507_FREM3, Exc_L4-5_RORB_FOLH1B) (Fig. 5b-d,h). *LAMC3* plays a role in cortical lamination in the mouse^35^ and mutations are implicated in human brain heterotopias and gyration defects^36,37^. Similarly, *S0X11*, a transcription factor that plays a role in embryonic and adult neurogenesis in the mouse brain^38^ and decreases in expression in the cerebral cortex during development^39,40^, was upregulated in paediatric Exc_L3-5_RORB_ESR1 and Exc_L2-3_LINC00507_FREM3 (Fig. 5b-c,h). *FNBP1L* (*TOCA-1*) was upregulated in Exc_L2-3_LINC00507_FREM3 and Exc_L2_LAMP5_LTK (Fig. 5c,e,h). FNBP1L promotes actin polymerisation, regulating neurite outgrowth, and declines in expression over the course of brain maturation in the rat^41^. Two genes, *STEAP2*, a metalloreductase, and the TNF receptor *TNFRSF25* (*DR3*), were higher in adult Exc_L3-5_RORB_ESR1 and Exc_L4-5_RORB_FOLH1B subtypes (Fig. 5b,d,h). *STEAP2* increases in expression during post-natal hippocampal maturation in mice^42^. *TNFRSF25* is activated post-natally in the mouse brain, where it may play a role in retention of motor control during aging^43^. These findings indicate that previously reported expression dynamics for these genes in mammalian models are conserved in the human temporal cortex. Importantly, our analysis reveals these patterns are specific to groups of excitatory neuron subtypes.

The majority of the DEGs were not shared across the cell types. For example, *FGF13* (*FHF2*) and *TENM1* were upregulated in paediatric Exc_L3-5_RORB_ESR1 (Fig. 5b). *FGF13* decreases in expression with age in the mouse brain, where it regulates post-natal neurogenesis^44^ and axonal formation^45^. *TENM1* is a member of the teneurin transmembrane protein family that regulates cytoskeletal organisation and neurite outgrowth, as well as shaping synaptic connections^46–48^. *KCNG1,* a voltage gated-potassium channel (Kv6.1), was upregulated in paediatric Exc_L2-3_LINC00507_FREM3 neurons (Fig. 5c), while *MYO16* (*MYR8*), an unconventional myosin protein, was upregulated in the Exc_L2_LAMP5_LTK subtype (Fig. 5e). Both of these genes decrease in expression with age in the mammalian brain^49,50^.

In line with our minimal marker analyses, fewer genes were differentially expressed in non-neuronal cells (Fig. 5f-g, Extended Data Table 10). In Astro_L1-6_FGFR3_SLC14A1, *PIK3R3* was upregulated in paediatric samples, while *PFKFB2* was downregulated. *PIK3R3* is involved in the PI3K-AKT growth signalling pathway, which is implicated in brain growth disorders^51^. *PFKFB2* is a bifunctional kinase/phosphatase that controls glycolysis. In contrast to our findings, PFKFB2 expression is higher in juvenile rat hippocampal astrocytes than in adults, where it may support energy demands during learning^52^. In oligodendrocytes, *NOTCH2* and *RRAS2* were both upregulated in paediatric samples. *Notch2* expression decreases in the rat cortex with age^53^ and is proposed to regulate glial differentiation^54^. These results provide new molecular candidates to expand our understanding of the mechanisms of astrocyte and oligodendrocyte maturation.

To explore the trajectories of DEG expression, we employed *psupertime* pseudotime trajectory analysis^55^, focussing on the four excitatory neuron sub-types with the highest number of DEGs. In support of our DESeq2 findings, several of the identified DEGs had non-zero psupertime coefficients and therefore represent genes that are relevant to the ordering of the cells in pseudotime^55^ (Fig. 5i; Exc_L3-5_RORB_ESR1: 13/47 [28%], Exc_L2-3_LINC00507_FREM3: 16/38 [42%], Exc_L4-5_RORB_FOLH1B 3/27 [11%] and Exc_L2_LAMP5_LTK: 5/18 [28%]; Extended Data Table 11). When considering the pseudotime trajectories for all DEGS in these excitatory neuron subtypes, the direction of the expression matched the DESeq2 results (Supplementary Fig. 2-5). The pseudotime trajectories revealed subtle expression dynamics within the analysed sample groups, showing that the majority of DEGs gradually increase in expression with age from childhood to adolescence, followed by a decrease in expression towards late adulthood.

Genes associated with intelligence quotient (IQ) and educational attainment (EA), as well as those associated with accelerated evolution in humans, have recently been shown to be enriched in adult temporal lobe cortical neurons, especially the Exc_L2-3_LINC00507_FREM3 subtype^56^. Since childhood is a key period of cognitive development^57^, we explored whether the same genes were found amongst our DEGs. Of the 149 DEGs found in at least one cell type, 20 (13.42%, p=0.02) are known to be significantly associated with EA^58^, 6 (4.02%, p=0.7) with IQ^59^ and 30 (20.13%, p=3.89E-07) with accelerated evolution in humans^60^. These included several genes that are upregulated in paediatric samples, such as *MYO16*, *KCNG1*, *FGF13* and *SOX11* (Extended Data Table 10).

Overall, we highlight several genes that are upregulated in children/adolescents which have known roles in brain development and have been associated with cognitive ability. Our analysis builds on previous knowledge by implicating specific cell subtypes and provides new candidate genes that likely contribute to cell type-specific maturation processes.

### Gene pathways involved in cellular respiration and synaptic functioning are enriched in paediatric cell types

We next used gene set enrichment analysis (GSEA) to conduct a broad analysis of the gene pathways that are differentially regulated across all brain cell types during brain maturation. 2,006 GOBP terms where enriched in the paediatric samples compared to the adults, while 866 were depleted (p<0.01 and q<0.1) (Extended Data Table 12). When focussing on the 25 most frequently enriched terms, the majority (10 terms) were associated with cellular respiration pathways (Fig. 6; Extended Data Table 12). Six were associated with intra-cellular transport, including transport of neurotransmitters, while five were linked to neurotransmitter release and synaptic plasticity. Three terms, including the top enriched term, were associated with protein translation and modification. The majority of depleted terms (10 terms) were associated with synaptic processes (Fig. 6). A further six depleted terms were connected to neuronal morphogenesis, including axon and dendrite morphogenesis. Two of the top depleted terms were associated with axon ensheathment. Interestingly, neither of these terms were significantly enriched in oligodendrocytes or OPCs, while they were associated with neuronal sub types, and microglia.

**Fig. 6:**
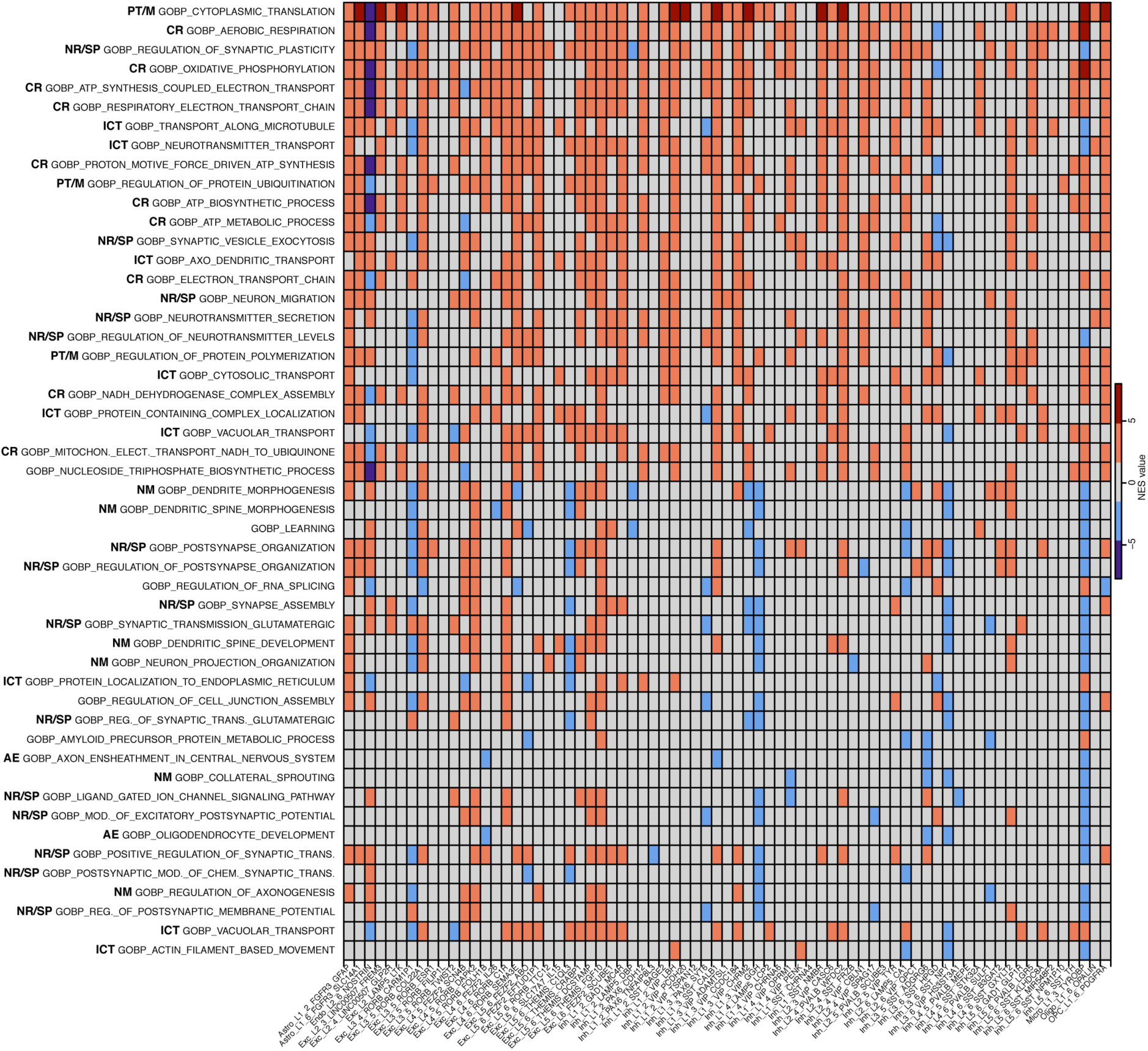
Pathways that are enriched or depleted across multiple cell types in paediatric samples. GSEA heatmap showing the top 25 most frequently enriched (top 25 rows) or depleted (bottom 25 rows) terms appearing across all cell types. Abbreviations in bold indicate the following categories as referred to in the text: AE, axon ensheathment; CR, cellular respiration; ICT, intracellular transport; NM, neuronal morphogenesis; NR/SP, neurotransmitter release/synaptic plasticity; PT/M, protein translation/modification. Only significantly (p < 0.01 and q < 0.1) terms are shown. NES value represents the normalized enrichment scores. Grey indicates that the term was not significantly enriched or depleted in the indicted cell type. See also Extended Data Table 12.

Overall, our GSEA analysis points towards putative genetic pathways that may drive maturation in the paediatric brain. Cellular respiration processes needed to support the higher metabolic rates in the brain during childhood^61^ may be enriched. Additionally, pathways related to strengthening synapses through neurotransmitter release may be enhanced. On the other hand, as synaptic pruning is underway^62^, pathways that promote synaptic growth may need to be suppressed.

### Cell type-specific expression of site-of-disease TBM biomarkers

The PCA aims to create reference atlases that can be used to improve our understanding of cell type-specific responses to disease in children^17^. Here, we used our snRNA-seq datasets to interrogate the cell type-specific expression of putative genetic biomarkers for TBM^23^. These biomarkers are enriched in the ventricular cerebrospinal fluid from children with TBM in comparison to controls with meningitis caused by other brain infections^23^.

66 of the 76 TBM biomarkers were expressed in our dataset, with similar expression across the two age groups, and genes clearly clustering according to their relative expression across the broad cell type categories (Extended Data Fig. 11). The genes with the highest relative expression in our data were expressed by non-neuronal cell types, which is in line with the view that immunological activity of supporting cells and their intercellular signalling interactions are important drivers of the immune response to TBM^63^. Several of these biomarkers (e.g. *FADS2*, *AMOT* and *ALDH6A1*) were enriched in the two astrocyte subtypes, potentially pointing towards a prominent role for astrocytes in the host response to TBM.

Our analyses also clearly revealed subsets of biomarkers that are more highly expressed by neuronal subtypes than non-neuronal cell types. This included biomarkers that were associated with Exc_L2_LAMP5_LTK, Exc_L2-4_LINC00507_GLP2R and Exc_L5-6_THEMIS_C1QL3 (*LYNX1*, *FAIM2*, *MAP1A*, *TUBB4A*). This is line with the finding that neuronal excitotoxicity is elevated in TBM^23^ and suggests that specific excitatory neuron subtypes may be contributing to this signal.

Interestingly, the two most enriched genes in the TBM biomarker dataset, *CXCL9* and *CXCL11*, were either completely absent from our datasets (*CXCL11*) or expressed by very few nuclei (*CXCL9*). The absence of these interferon-inducible chemokines in our datasets from uninfected tissue, supports the proposition that they are indeed biomarkers from the site-of-disease^64^ in both adults and children with TBM, and could also reflect the contribution of peripheral immune cells recruited to the brain during infection.

## Discussion

The brain is the most complex organ in the human body, which continuously changes as we mature. Here, we begin to unmask the molecular mechanisms guiding these processes in the temporal cortex, using single cell and spatial transcriptomics to compare similar cell types between paediatric and adult datasets.

To facilitate accurate comparisons of cell types across age groups, we used the existing Allen Brain Map MTG cell atlas^1^ to annotate our datasets. This demonstrated that the reference atlas, generated from adult snRNA-seq datasets, is indeed generalisable^31^, and can be used to classify cell types from samples of different ages. This generalisability is essential for healthy human reference atlases to serve as a baseline to improve our understanding of human development and disease^3^. Our samples and those in the reference MTG cell atlas include neurosurgical tissue from donors with epilepsy, and while the analysed tissue is not from the site of pathology, it is important to view our findings in light of the patient diagnosis. Previous research comparing gene expression between the neurosurgical and post-mortem samples used in the reference MTG cell atlas found a strong correlation of expression between cell types across conditions^1^. In addition, a comparison of samples from 45 adult donors with epilepsy to the post-mortem samples from the reference MTG cell atlas found a similar number of genes and similar cell abundance per cell subclass across tissue sources, however they did find more variation for these parameters in neurosurgical samples^65^. As more paediatric MTG samples of post-mortem and neurosurgical origin become available, it will be important to conduct similar analyses to determine if these findings hold for the paediatric temporal cortex.

Our machine-learning marker gene analysis shows that while the cell type classifications, which are based on the expression of thousands of genes, can be transferred onto new datasets, the minimal markers that define the cell types do vary across datasets. Only a quarter of our *NS-Forest* minimal markers overlap with the existing MTG cell atlas minimal markers^29^. The differences in the single cell transcriptomics technologies used to generate our dataset and the MTG cell atlas may account for much of this discrepancy. Nonetheless, our analyses suggest that some of our markers may provide better discrimination between cell types than existing markers. These results highlight a challenge that the HCA faces to revise cell type markers as more datasets are made available to ensure that the cell type classification is as widely applicable as possible.

Similar to analyses of aging in the mouse^66^, our analyses show there is little change in cell type composition within the temporal cortex during human brain maturation. However, our differential expression analysis highlights differences in cell states between specific paediatric and adult cell subtypes. Recently, the supragranular excitatory pyramidal neurons in the MTG have been shown to have high transcriptional diversity^1,67^, large arborisations^68^ and electrophysiological properties that impact signal integration and encoding^69–72^ in ways that may contribute to cognition. Since cognitive ability is a key feature that is established during childhood^68^, our analysis offers an opportunity to explore how cell type-specific gene expression dynamics contribute to cognitive development. Interestingly, two of the 21 highlighted cell types were the layer 2/3 excitatory neurons, Exc_L2_LAMP5_LTK and Exc_L2-3_LINC00507_FREM3, that have recently been associated with human cognition^56^. In line with these findings, several of the DEGS associated with these cell types, including *FNBP1L*^73^ and *SOX11*^74^, have been implicated in cognitive ability and intelligence. Overall, our data points towards genes that may play roles in cognitive development specifically within these excitatory neurons.

The relatively low number of genes implicated in our differential expression analysis in comparison to similar studies in mouse^66^ suggests that the difference between the paediatric and adult brain are subtle. However, the inherent high variability in human gene expression data may mask some of the differential gene expression in our limited sample. Nonetheless, our pseudotime trajectory analyses reveals some of the expression dynamics that may be occurring during childhood, with many genes rising in expression towards adolescence and dropping off in adulthood. As the HCA database for the human temporal cortex expands, it will be important to build on these analyses with more samples. Binning of samples of similar age will provide a higher resolution analysis of cell type-specific gene expression trajectories over the course of brain maturation.

Finally, we have provided the first single nucleus gene expression datasets for the brain that includes data from black Southern African donors, thus increasing the diversity of the HCA database. We demonstrate how this resource can be used to deconvolute site-of-disease biomarker analyses for TBM, pinpointing which cell types may be driving altered gene expression profiles in the brain. Importantly, these investigations have the potential to contribute to the development of effective treatments, that are tailored to specific needs of both adult and paediatric patients.

## Methods

### Human samples

Ethical approval was granted for the collection and use of paediatric and adult human brain tissue by the University of Cape Town Human Research Ethics Committee (UCT HREC REF 016/2018; sub-studies 146/2022 and 147/2022). The human brain tissue samples used to generate new datasets were obtained by informed consent for studies during temporal lobe surgical resections to treat epilepsy and/or cancer performed at the Red Cross War Memorial Children’s Hospital and Constantiaberg Mediclinic in Cape Town, South Africa. The samples used in this study were of temporal cortex origin and represent radiologically and macroscopically normal neocortex within the pathological context (details in Extended Data Table 1). Race was recorded by the clinical teams based on their knowledge of the donors. The category “black South African” includes both black and mixed race ancestries. Upon resection, samples were placed in carbogenated ice-cold artificial cerebral spinal fluid (aCSF) containing in (mM): 110 choline chloride, 26 NaHCO_3_, 10 D-glucose, 11.6 sodium ascorbate, 7 MgCl_2_, 3.1 sodium pyruvate, 2.5 KCl, 1.25 NaH_2_PO_4_, and 0.5 CaCl_2_ (300 mOsm) and immediately transported to the laboratory (∼20 minutes). Tissue blocks containing the full span from pia to white matter were prepared and either flash frozen in liquid nitrogen or embedded in optimal cutting temperature compound (OCT) and stored at -80°C. The OCT-embedded samples were flash frozen in a 10×10 mm^2^ cryomold which was either frozen directly in liquid nitrogen or placed in a container of isopentane (Merck) which was in turn placed in liquid nitrogen at the same level as the isopentane. The publicly available snRNA-seq datasets^24^, generated from samples obtained during elective surgeries performed at Universitair Ziekenhuis Leuven, Belgium, were downloaded from the Sequence Read Archive database.

### Nuclei isolation for snRNA-seq

Nuclei were isolated according to a protocol adapted from Habib et al. (2017)^75^ and the 10X Genomics nuclei isolation protocol (CG000124, User Guide Rev E). Frozen brain tissue was homogenised in a dounce-homogeniser containing 2 ml ice-cold lysis solution (Nuclei EZ Lysis Buffer [Sigma-Aldrich, NUC101] or Nuclei PURE Lysis buffer [Sigma-Aldrich, NUC201] with 1 mM dithiothreitol [DTT, Promega, P1171, US] and 0.1% Triton X-100 [Sigma-Aldrich, NUC201-1KT, US]). Homogenisation was done 20 times with the loose pestle A followed by 20 times with the tight pestle B. An additional 2 ml lysis solution was added, and the sample was incubated for 5 minutes on ice. The sample was centrifuged at 500 x g for 5 minutes at 4°C after which the supernatant was discarded and the nuclei resuspended in 3 ml ice cold nuclei suspension buffer (1x phosphate-buffered saline [PBS, Sigma-Aldrich, P4417-50TAB, US]), 0.01% bovine serum albumin [BSA, Sigma-Aldrich, A2153-10G, US], and 0.2 U/µl RNAsin Plus RNase inhibitor [Promega, N2615, US]). Resuspended nuclei were passed through a 40 µm filter and centrifuged at 900 x g for 10 minutes at 4°C. The supernatant was discarded and pelleted nuclei were resuspended in 3 ml blocking buffer (1xPBS [Sigma-Aldrich, P4417-50TAB, US], 1% BSA [Sigma-Aldrich, A2153-10G, US], 0.2 U/µl RNAsin Plus RNase inhibitor [Promega, N2615, US]).

To remove myelin debris, 30 µl of myelin removal beads [Miltenyi Biotec. 130-096-733, US] was added to the solution which was mixed by gently pipetting 5 times. The sample was incubated for 15 minutes at 4°C after which it was mixed with 3 ml blocking buffer and centrifuged at 300 x g for 5 minutes at 4°C. The supernatant was removed and the nuclei were resuspended in 2 ml clean blocking buffer. The sample was transferred to a 2 ml tube and placed on a Dynamag magnet for 15 minutes at 4°C. The supernatant was transferred to a new tube and stored on ice. An aliquot of trypan blue stained nuclei was counted using a haemocytometer to determine the nuclei concentration and the volume to use in snRNA-seq library preparation.

### 10X Genomics snRNA-seq library preparation

snRNA-seq library preparation was carried out using the 10x Genomics Chromium Next Gem Single Cell 3’ Reagent Kit (v3.1) according to manufacturer’s protocols (CG000204, User Guide Rev D), targeting 10,000 nuclei per sample. At step 2.2d and 3.5e, the libraries were amplified using 11 cycles and 13 cycles, respectively. Library quality and concentration was assessed using either the TapeStation or Bioanalyser (Agilent) and Qubit (Invitrogen) at the Central Analytical Facility (CAF, University of Stellenbosch). cDNA libraries were sequenced by Novogene (Singapore) on either the Illumina HiSeq or NovaSeq system using the Illumina High Output kits (150 cycles).

### snRNA-seq read alignment and gene expression quantification

Fastq files were aligned to the human reference transcriptome (GRCh38) and quantified using the count function from the 10X Genomics Cell Ranger v6.1.1 software (Cell Ranger, RRID SCR_017344) (Code availability: script 1). The inclusion of introns was specified in the count function. An automatic filtering process was performed to remove barcodes corresponding to background noise which have very low UMI counts.

### snRNA-seq quality control

The resulting count matrices were processed using a pipeline adapted from the Harvard Chan Bioinformatics Core (https://hbctraining.github.io/scRNA-seq_online/). The filtered gene barcode matrix for each sample was imported into R (V.4.2.0) using the Read10X function from the Seurat (v.2.0)^25^. Nuclei-level filtering was performed to remove poor quality nuclei according to their number of UMIs (nUMIs) detected, number of genes detected (nGene), number of genes detected per UMI (log10GenesPerUMI), and the fraction of mitochondrial read counts to total read counts (mitoRatio) (Code availability: script 2). Nuclei that met the following criteria were retained: nUMI > 500, nGene > 250, log10GenesPerUMI > 0.8 and mitoRatio < 0.2. Gene-level filtering was performed to remove genes that had zero counts in all nuclei, remove genes expressed in fewer than 10 nuclei, and remove mitochondrial genes from the gene by cell counts matrix. Three doublet removal tools namely DoubletFinder^76^(Code availability: script 3), DoubletDecon^77^ (Code availability: script 4), and Scrublet^77^ (Code availability: script 5,6) were used to identify doublets for each dataset individually. The sample-specific parameters of each of the tools were adjusted according to the specified guidelines. To achieve a balance between the false positive and false negative rate of the different doublet detection tools, all doublets identified by DoubletFinder as well as the intersection of the doublets identified by DoubletDecon and Scrublet, were removed^77^.

### snRNA-seq data normalization, integration and clustering

Principal component analysis was performed to evaluate known sources of within-sample variation between nuclei, namely the mitoRatio and cell cycle phase (Code availability: script 7). The UMI counts of the 3000 most variable features were normalised and scaled on a per sample basis by applying Seurat’s SCTransform function with mitoRatio regressed out. A Uniform Manifold Approximation and Projection (UMAP) analysis was performed on the merged object to assess whether integration was necessary. The datasets were subsequently integrated using Seurat’s SelectIntegrationFeatures, PrepSCTIntegration, FindIntegrationAnchors, and IntegrateData functions (Code availability: script 7). To cluster the datasets following integration, dimensionality reduction was first performed using UMAP embedding, specifying 40 dimensions (Code availability: script 8). The Seurat FindClusters function was then applied at a resolution of 0.8.

### snRNA-seq cluster annotation

Two levels of annotation were performed. Clusters were initially annotated as one of the major brain cell types (level 1 annotation) based on the expression of known markers genes (Code availability: script 9). Label transfer was then performed using Seurat’s TransferData function with Allen Brain Map MTG atlas^1^ as a reference dataset (level 2 annotation) (Code availability: scripts 10-11). This resulted in each barcode in the query dataset receiving a predicted annotation based on a similarity score to an annotated cell type in the reference. Barcodes were then filtered to remove those with discordant level 1 and level 2 annotations (e.g. barcodes with “oligodendrocyte” level 1 annotation and “Exc_L4-5_RORB_FOLH1” level 2 annotawon) (Code availability: script 12). To validate the annotation, the expression of known marker genes was assessed. Cosine similarity scores were computed to compare the transcriptomic similarity of each of the annotated query cell types to the 75 reference MTG cell types using the SCP package (https://github.com/zhanghao-njmu/SCP) (Code availability: script 13). This was achieved by computing cosine similarity scores for each pair of query and reference cell types using the expression of the top 2000 shared highly variable features between the query and reference datasets. The log normalised expression counts were used for this purpose (RNA assay, data slot). To assess the difference between the paediatric and adult datasets relative to the reference, the above cosine similarity analysis was repeated on the paediatric and adult datasets individually (Code availability: script 13).

### NS-Forest machine learning marker analysis of snRNA-seq datasets

The NS-Forest tool (v2.0)^29,30^ was used to identify combinations of marker genes uniquely defining each annotated cell type (Code availability: script 14-15) in the paediatric and adult datasets separately. The number of nuclei per sample was randomly down-sampled to that of the sample with the fewest nuclei (n=4,865). A random-forest model was used to select a maximum of 15 marker genes per cell type based on them being both highly expressed as well as uniquely expressed within a cell type compared to other cell types (i.e., the top Gini Index ranked features with positive expression values). The number of trees chosen for this model was 30,000, the cluster median expression threshold was set to the default value of zero, the number of genes used to rank permutations of genes by their F-beta-score was 6, and the beta weight of the F score was set to 0.5. The aforementioned parameters were set according to the parameters described in Aevermann et al. (2021)^29^, allowing the outputs to be directly compared to their markers and to the Allen Brain Map MTG atlas minimal markers^1^. To assess the relevance of these markers in terms of their capacity to distinguish different cell types in a UMAP analysis, the SCT and integration methods were repeated using either a random set of genes or the NS-Forest markers as anchors^29^ (Code availability: script 16).

### DESeq2 age-dependent differential gene expression analysis of snRNA-seq datasets

DESeq2^33^ was used to identify genes that were differentially expressed with age (Code availability: script 17). The unnormalized counts were aggregated across all nuclei for each cluster and sample to generate a ‘pseudobulk’ counts matrix with the counts from technical replicates collapsed to the level of biological replicates. Genes were filtered to only include those expressed in more than 10% of nuclei for a given cell type. Principal component analysis was performed on each cell type separately in order to assess the variation between samples and determine which variables were contributing most to inter-sample variation from a set of possible variables. The collapsed counts served as input into DESeq2’s DESeqDataSetFromMatrix function in which the design formula ∼single_cell_chemistry + age_group was specified to treat the age_group (paediatric vs adult) as the variable of interest while the effect of single_cell_chemistry (version2 vs version3 chemistry) was regressed out. A hypothesis test was performed using the Wald test. The null hypothesis for each gene was that there is no difference in gene expression between the sample groups (i.e Log2Fold Change = 0). A Wald test statistic was determined for each gene together with the associated p-value after which the p-values were adjusted for multiple testing using the Benjamini-Hochberg method. Positive log2 Fold Changes represent genes which are upregulated in paediatric samples compared to adult samples (p_adj_ <0.05).

### Pseudotime trajectory analysis with psupertime

To validate the differentially expressed genes identified with DESeq2, a pseudotime trajectory analysis was performed for a subset of excitatory neuron subtypes using the psupertime package^55^ (Code availability: script 18). Psupertime is a supervised approach that uses time-series labels as input to improve the identification of time-varying genes. Each cell type of interest (Exc L3-5 RORB ESR1, Exc L2 LAMP5 LTK, Exc L4-5 RORB FOLH1B, Exc L2-3 LINC00507 FREM3) was individually sub-setted from the Seurat object after which a single cell experiment (sce) object was generated using the log normalized counts (RNA assay, data slot). The donor age (4, 5, 7, 9, 15, 20, 24, 26, 31, 41, 50) was included as metadata in the object. The psupertime function was applied to the sce object with the sel_genes argument specifying all genes to be used. An automatic filtering step was performed to remove genes expressed in fewer than 10% of cells for each cell type. As an output of the function, the beta coefficients for the association of each gene with pseudotime were extracted and plots were generated showing the expression trajectories of the DESeq2 DEGs with pseudotime. Additionally, the overlap between the DESeq2 DEGs and genes changing as a function of pseudotime (Psupertime-relevant genes) was determined.

### Pathway enrichment analysis of snRNA-seq datasets

GO analysis of NS-Forest marker genes was performed on the gProfiler web server^78^ using default settings (p_adj_ <0.05) with “highlight diver terms in GO” selected.

DEGs identified by DESeq2 (see Extended Data Table 10) that were associated with EA and IQ, as well as those associated with accelerated evolution in humans (HARs), were determined by comparing the list of neuronal DEGS to the EA, IQ and HAR gene lists used by Driessens et al. (2023)^56^, which were subsets of the lists from Lee et al. (2018)^58^, Savage et al. (2018)^59^ and Doan et al. (2016)^60^ respectively. A hypergeometric test was performed to test the significance of the results relative to chance (Code availability: script 19).

GSEA on the DESeq2 output for all genes was performed using the Broad Institute’s GSEA software (https://www.gsea-msigdb.org/gsea/msigdb) (Code availability: script 20). GSEA aggregates the information from many genes to identify enriched functional pathways, allowing us to interrogate the gene signature changes across all cell types, including those that did not show any significant DEGs^66^. The gene lists for each cell type were queried against the C5 GO Biological Processes collection comprising of gene sets derived from the GO Biological Process ontology. The input lists of genes were ranked according to the -log(p-value)*log_2_FoldChange for each gene. The parameters specified to the GSEA function included number of permutations (nperm)=1000, minimum gene set size (set_min=15), maximum gene set size (set_max=200), excludes genes that have no gene symbols (collapse)= No_Collapse, value to use for the single identifier that will represent all identifiers for the gene (mode)=Max_probe, normalised enrichment score method (norm)= meandiv, weighted scoring scheme (scoring_scheme) = classic. Positive Normalised Enrichment Scores (NES) represent genes that were upregulated in the paediatric population compared to the adult population (p<0.01 and q<0.1). To visualise the output of universally enriched pathways across multiple cell types, the top 25 most frequently appearing positively and negatively associated terms were plotted. Additionally, for five cell types of interest [which had DEGs meeting the threshold of p<0.05 and abs(log_2_FC)>0.1], the top 5 positively associated terms were plotted.

### Analysis of site-of-disease TBM markers

The dittoheatmap function from the dittoSeq package^79^ was used to generate heatmaps for the expression of the TBM biomarkers (upregulated genes listed in Rohlwink et al. 2019, Supplementary Table 5^23^) across cell types in the paediatric and adult datasets individually. Additionally, Seurat’s dotplot function^25^ was used to visualize the level of expression and proportion of nuclei expressing the markers across cell types (Code availability: script 21). Prior to generating the plots, the TBM marker genes were filtered to remove those expressed in 15 nuclei or fewer across all cell types. Gene counts for each marker were aggregated across cell types and scaled. The markers were clustered according to their expression profiles using dittoheatmap’s default hierarchical clustering method (Euclidean, complete). The clustering order and dendrogram from this output for the peaditaric datasets were used to generate dotplots for both peaditaric and adult datasets (Code availability: script 21).

### snRNA-seq data plots

Plots were produced with Seurat^25^, ggplot2^80^, ShinyCell^81^ and Microsoft Excel.

### 10x Genomics Visium library preparation

Frozen OCT embedded temporal cortex tissue samples were scored using a pre-chilled razor blade to fit in the Spatial Gene Expression slide capture areas. 10 μm-thick sections were cut using a cryostat (Leica CM1860/CM1950) and collected onto the Spatial Gene Expression slide capture areas. Two replicate sections of the 15-year-old (10 μm apart) and two replicate sections of 31-year-old (40 μm apart) were collected. The spatial Gene Expression slides with tissue sections were stored in a sealed container at -80°C. Captured sections were Haematoxylin and Eosin (H&E) stained according to the 10x Genomics Demonstrated Protocol Guide (CG000160, Rev B). Brightfield images of the stained sections were captured using an EVOS M5000 microscope (Thermo Fisher Scientific) at 20x magnification without coverslipping. Overlapping images of the sections including the fiducial frame were stitched together using Image Composite Editor-2.0.3 (Microsoft). Visium libraries were prepared from the stained tissue sections following the Visium Spatial Gene Expression Reagents Kit User Guide (CG000239, Rev D). At Step 1.1 the tissue was permeabilised for 12 minutes as determined using the Visium Spatial Gene Expression Tissue Optimisation User Guide (CG000238, Rev D). At Step 3.2, cDNA was amplified using 20 cycles. Library quality and concentration was assessed using TapeStation (Agilent) and Qubit (Invitrogen) at the Central Analytical Facility (CAF, University of Stellenbosch). Libraries were sequenced by Novogene (Singapore) on the Illumina NovaSeq system using the Illumina High Output kits (150 cycles).

### Visium read alignment and gene expression quantification

The H&E images were processed using the 10X Genomics Loupe Browser V4.0 Visium Manual Alignment Wizard. 10X Genomics Space Ranger *count* (10X Space Ranger V1.3.0) was used to perform alignment of FASTQ files to the human reference transcriptome (GRCh38), tissue detection, fiducial detection and barcode/UMI counting.

### cell2location analysis of Visium datasets

The average number of nuclei per Visium spot was determined using Vistoseg^82^ (Code availability: script 22). Cell2location (version 0.7a0)^26^ was used to spatially map the brain cell types by integrating the Visium data count matrices (Space Ranger output) with the annotated snRNAseq datasets (Code availability: script 23). To avoid mapping artifacts, mitochondrial genes were removed from the Visium datasets prior to spatial mapping. Reference signatures of the 75 annotated cell populations were derived using a negative binomial regression model using the default values (Code availability: script 24). Unnormalized and untransformed snRNA-seq mRNA counts were used as input in the regression model for estimating the reference signatures (Code availability: script 24. The snRNA-seq mRNA counts were filtered to 14,209 genes and 144,438 cells. The cell2location model for estimating the spatial abundance of cell populations was filtered to 14,197 genes and 14,324 cells that were shared in both the snRNA-seq and Visium data. The following cell2location parameters were used: training iterations = 30,000 cell per location, N^ = 7 (estimated using Vistoseg segmentation results), Normalization (ys) alpha prior = 20 (Code availability: script 25). To visualise the cell abundance in spatial coordinates 5 % quantile of the posterior distribution was used, which represents the value of cell abundance that the model has high confidence in (Code availability: script 26). Cell2location’s Non-negative Matrix Factorization (NMF) was used to identify cellular compartments and cell types that co-locate from the cell type abundance estimates. NMF was tested using a range of factors (5 to 30) for the “n_fact” parameter (Code availability: script 26). n_fact=15 was chosen as it clearly grouped the oligodendrocyte, astrocyte and excitatory neuron cell sub-types into known tissue zones i.e. the layers of the cortex (Code availability: script 27).

### BayesSpace analysis of Visium datasets

The raw gene expression counts from Space Ranger were normalized, log transformed and principal component analysis was performed on the top 2000 highly variable genes. To obtain high-resolution gene expression for selected genes, the principal component values were mapped back to their original log-transformed gene expression space (spot level) using the default BayesSpace^34^ regression (Code availability: script 28). To do this the principal components from the original data were used as predictors in training the model for each gene, in which the results were the measured gene expression at the spot level. The trained model was then used to predict the gene expression at sub spot level using high resolution PCs. The high-resolution model was trained using default values except for the following parameters: 7 PCs, Number of clusters = 8, nrep = 100,000, burn-in = 10,000. The BayesSpace outputs for each sample were quantified for spots with expression level > 0 and displayed as boxplots (Code availability: script 29).

### *In situ* Hybridisation Chain Reaction (HCR) on frozen human tissue sections

10 μm thick frozen sections were collected on Histobond+ slides (Marienfeld) and stored at - 20°C. The *In situ* HCR protocol was carried out on tissue sections as detailed in Choi et al. (2016)^83^ using reagents, probes and hairpins purchased from Molecular Instruments. Probes were ordered for the following genes: *RELN* (NM_005045.4), *FABP7* (CR457057.1), *AQP4* (NM_001650.5), *RORB* (NM_006914.4), *CLSTN2* (NM_022131.3) and *TSHZ2* (NM_173485.6). When necessary to quench lipofuscin autofluorescence, sections were rinsed after HCR in 1x PBS and treated with 200 μl TrueBlack (Biotium) for 30 sec. Slides were rinsed in PBS, stained with Hoescht (Thermofisher) and mounted using SlowFade Gold Antifade Reagent (Invitrogen). Sections were imaged using the LSM 880 Airyscan confocal microscope (Carl Zeiss, ZEN SP 2 software) using the 40X or 60X objective.

### MERFISH analysis on frozen temporal cortex tissue sections

10 μm thick frozen sections were cut from frozen OCT embedded temporal cortex tissue samples using a cryostat (Leica CM1950). Sections from a peaditaric and adult sample were collected onto the same MERSCOPE coverslip (VIZGEN 2040003), fixed and stored in 70% ethanol following the instructions in the VIZGEN protocol (Fresh & Fixed Frozen Tissue Sectioning & Shipping Procedure Rev A, Doc. number 91600107). The slide was processed on the VIZGEN MERSCOPE system by the MRC Weatherall Institute of Molecular Medicine Single Cell Facility (University of Oxford) within 1 month of storage. Sections were photobleached for 10 hours at 4°C and then washed in 5 ml Sample Prep Wash Buffer (VIZGEN 20300001) in a 5 cm petri dish. Sections were incubated in 5 ml Formamide Wash Buffer (VIZGEN 20300002) at 37°C for 30 min and hybridized at 37°C for 36 to 48 hours by using 50 μl of VIZGEN-supplied custom Gene Panel Mix according to the manufacturer’s instructions. Following hybridization, sections were washed twice in 5 ml Formamide Wash Buffer for 30 min at 47°C. Sections were then embedded in acrylamide by polymerizing VIZGEN Embedding Premix (VIZGEN 20300004) according to the manufacturer’s instructions. Following embedding, sections were digested in Digestion Pre-Mix (VIZGEN 20300005) and RNase inhibitor (New England Biolabs M0314L) for 3 h at 37°C and then cleared for 16 to 24 hours with a mixture of VIZGEN Clearing Solution (VIZGEN 20300003) and Proteinase K (New England Biolabs P8107S) according to the Manufacturer’s instructions. Following clearing, sections were washed twice for 5 min in Sample Prep Wash Buffer (PN 20300001) and then stained with VIZGEN DAPI and PolyT Stain (PN 20300021) for 15 min followed by a 10 min wash in Formamide Wash Buffer. Formamide Wash Buffer was removed and sections were washed with Sample Prep Wash Buffer during MERSCOPE imaging set up. A mixture of 100 ml of RNAse Inhibitor (New England BioLabs M0314L) and 250 ml of Imaging Buffer Activator (PN 203000015) was added to the cartridge activation port to a prethawed and mixed MERSCOPE Imaging cartridge (VIZGEN PN1040004). 15 ml mineral oil (Millipore-Sigma m5904-6X500ML) was added on top of the activation port and the MERSCOPE fluidics system was primed according to VIZGEN instructions. The flow chamber was assembled with the section coverslip according to VIZGEN specifications and the imaging session was initiated after collection of a 10X mosaic DAPI image and selection of the 1cm^2^ imaging area. MERFISH data was visualised using the VIZGEN MERSCOPE Vizualizer software (version 2.3.3330.0).

## Data Availability

All scripts used to analyse the data are indicated in the methods section and are available in the supplementary material. A description of the raw and analysed data files will be made available on the University of Cape Town’s <u>ZivaHub</u> data sharing platform on publication. As the data is from living donors, access to the data will be mediated through contact with the corresponding author. A ShinyApp will be made publicly available on publication for exploration of the annotated snRNA-seq data.

## Acknowledgments

This research was support by a Royal Society/Global Challenges Research Fund/African Academy of Sciences Future Leaders African Independent Researcher (FLAIR) Fellowship to D.H. (FLR\R1\191008), a Royal Society/Global Challenges Research Fund FLAIR Collaboration grant to T.S.S. and D.H. (FCG\R1\201023), a National Research Foundation (NRF) Research Development Grant for Y-rated Researchers Award to DH (CSRP210415595025), a University of Cape Town (UCT) Building Research Active Academic Staff Grant award to D.H. and a National Institutes of Health (NIH) R21 Exploratory/Development Grant to D.H., J.V.R., C.G.D. and M.J. (TW011225). C.S. was supported by a Harry Crossley Research Scholarship, an Oppenheimer Memorial Trust scholarship, an NRF scholarship and a UCT Vice Chancellor’s Research Scholarship. S.F. was supported by a DAAD-NRF joint In-country Scholarship and a UCT Vice Chancellor’s Research Scholarship. M.B.V. was supported by an EMBO long-term fellowship (ALTF 415-2018) and a Claude Leon Foundation research fellowship. A.F. was supported by the NRF SARChI Chair of Clinical Neurosciences. Computations were performed using facilities provided by the University of Cape Town’s ICTS High Performance Computing team: <u>hpc.uct.ac.za</u>. This publication is part of the Human Cell Atlas: www.humancellatlas.org/publications/

## Contributions

C.S. and S.F. conducted the snRNA-seq experiments. C.S. conducted the majority of snRNA-seq analyses. R.M. conducted the Visium and HCR experiments and analysis. J.M. conducted additional HCR experiments. S.Q. provided additional bioinformatics support. M.B.V. liaised with neurosurgeons and prepared all neurosurgical brain tissue samples. R.M., J.B. and J.M.N.E conducted the neurosurgeries and provided donor metadata. C.S., T.S.S., M.G. and D.H. conceptualised, conducted and analysed the MERFISH experiments. U.K.R., M.Z., J.V.R, C.G.D, A.F, and D.H. conceptualised the study and raised funds. D.H., C.S. and R.M wrote the manuscript. D.H. conducted additional analyses and supervised the project. All authors read and commented on the manuscript.

---

**Extended Data Fig. 1:**
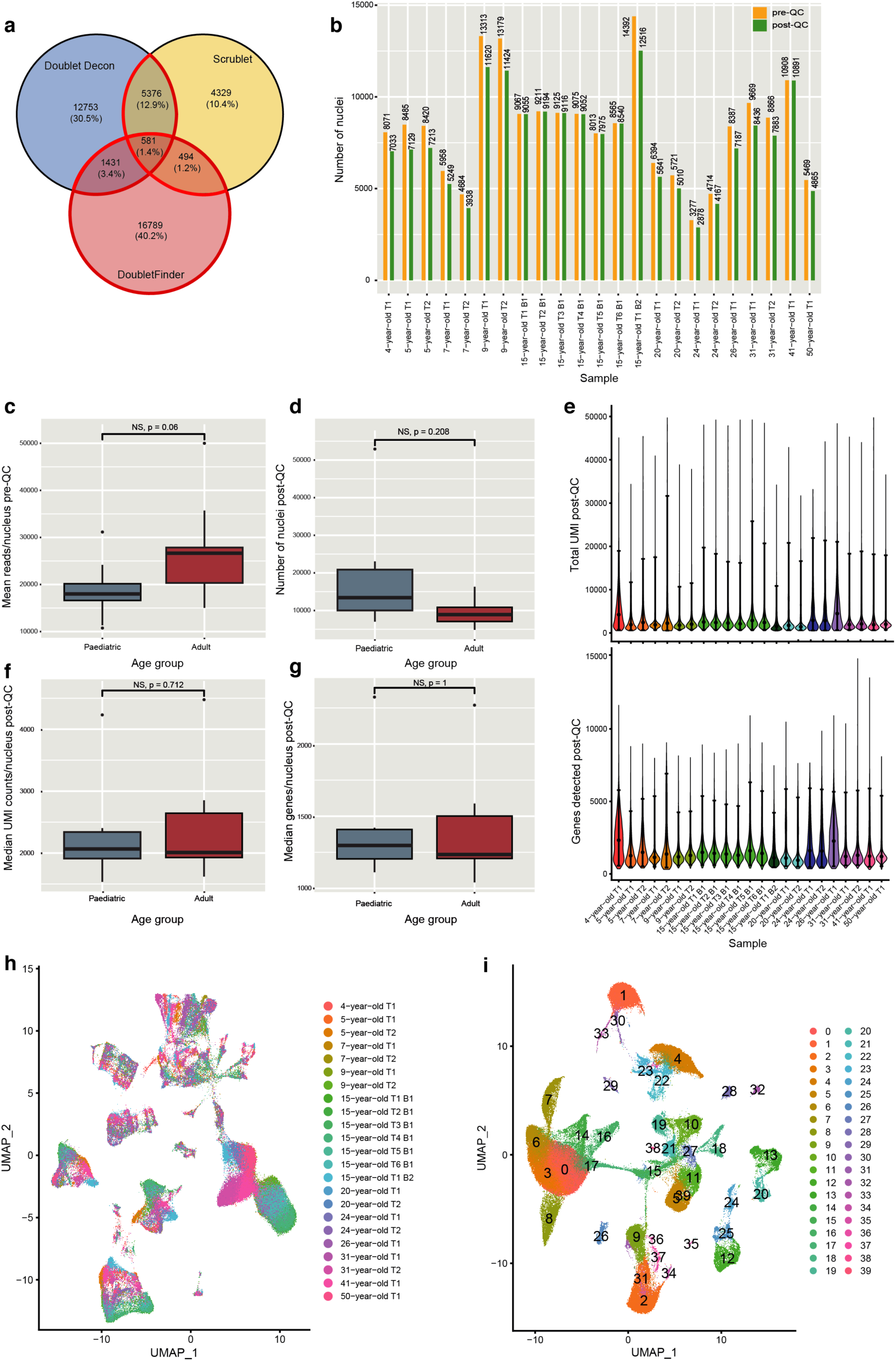
Nuclei quality control (QC) and clustering. **a**, Number of doublets identified across all 23 datasets by DoubletDecon, DoubletFinder, and Scrublet. Red outline indicates the subset of barcodes called as doublets that were removed. **b**, Total number of nuclei per dataset before (yellow) and after (green) QC. **c**, Mean number of reads per nucleus (y axis) by dataset before QC split by age group (x axis). p value determined by two-tailed Welch’s t-test. **d**, Number of nuclei (y axis) by sample after QC split by age group (x axis). p value determined by Brunnermunzel permutation test**. e**, Violin plots showing the number of unique molecular identifiers (UMIs) (top) and the number of genes detected (bottom) per nucleus per sample after QC. Black dots indicate the median value. Error bars show 95% confidence intervals. **f**,**g**, Median number of UMIs (2,263 paediatric and 2,011 adult) (**f**) and the median number of genes (1,372 paediatric and 1,226 adult) (**g**) detected per nucleus (y axes) by sample after QC split by age group (x axis). p values determined by two-tailed Brunnermunzel permutation test. **h**, UMAP plot for the 23 datasets prior to integration. **i**, UMAP plot showing the resulting clusters determined by the shared nearest neighbour algorithm. Data in all box plots represent mean ± sem for six paediatric and six adult samples. No significant differences were detected between paediatric and adult samples. B, biological replicate; NS, not significant; T, technical replicate. See also Extended Data Table 2.

**Extended Data Fig. 2:**
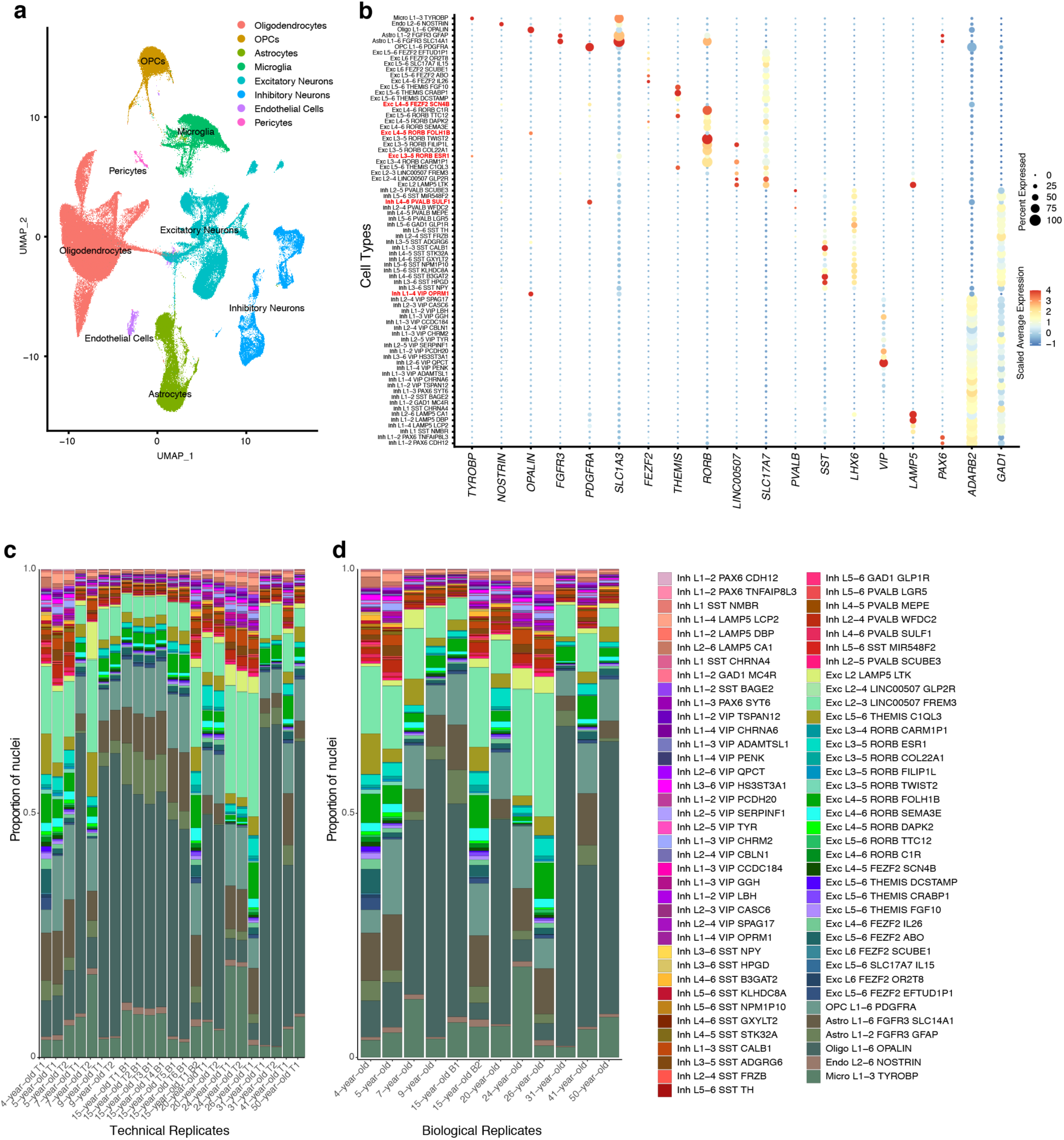
Annotation and assessment of cell composition across datasets. **a**, UMAP plot showing cluster annotation at the level of major brain cell types (level 1 annotation). **b**, Examination of known cell type-specific marker genes (x axis) after label transfer classify each nucleus according to the Allen Brain Map MTG atlas^1^ (level 2 annotation) (y axis) (left). Off-target gene expression is evident in several cell types (marked in red), which is likely due to multiplets or nuclei contaminated with ambient mRNA. **c-d**, Stacked barplots after filtering to retain nuclei with high confidence annotations showing the proportion of nuclei per cell type (y axis) for each technical replicate (**c**) or biological replicate (**d**) (x axis) out of the total number of nuclei for each group. Samples with technical replicates showed high degrees of similarity in cell composition between their replicates (**c**). Technical replicates from each donor were merged to allow comparisons between the 12 samples (**d**).

**Extended Data Fig. 3:**
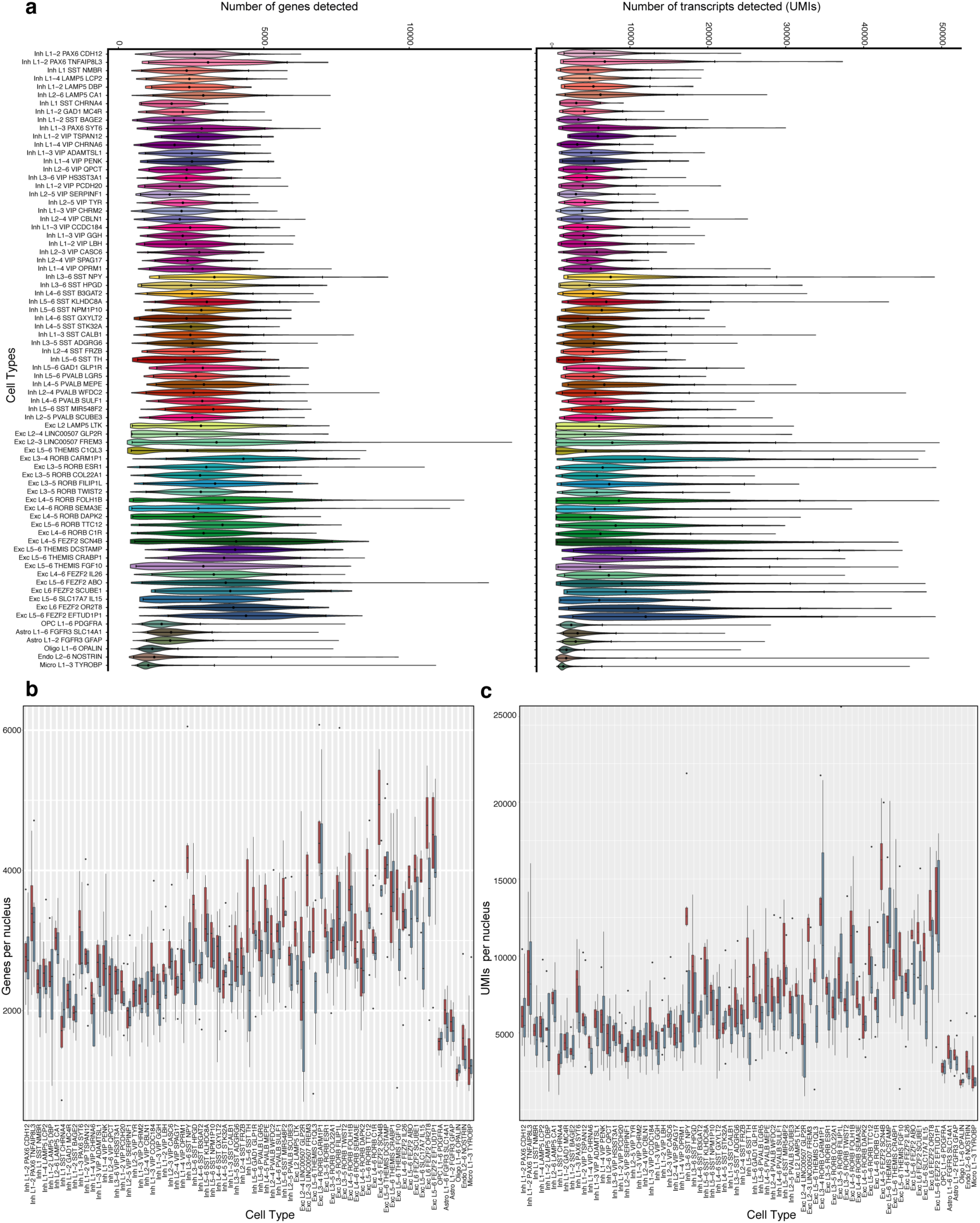
Assessment of the sequencing metrics for the annotated cell types. **a**, Violin plots showing the distribution of the number of genes (left) and transcripts (right) detected per nucleus per cell type across all datasets. Black dots indicate the median value. Error bars show 95% confidence intervals. **b**,**c**, Boxplots showing the number of genes (**b**) and the number of UMIs (**c**) (y axis) detected per cell type per sample (x axis) split by age group (red: adult, grey: paediatric). Data in all box plots represent mean ± sem for six paediatric and six adult samples for each cell type. No significant differences were detected (i.e. padj > 0.05). See Extended Data Table 3 for details of statistical tests performed.

**Extended Data Fig. 4:**
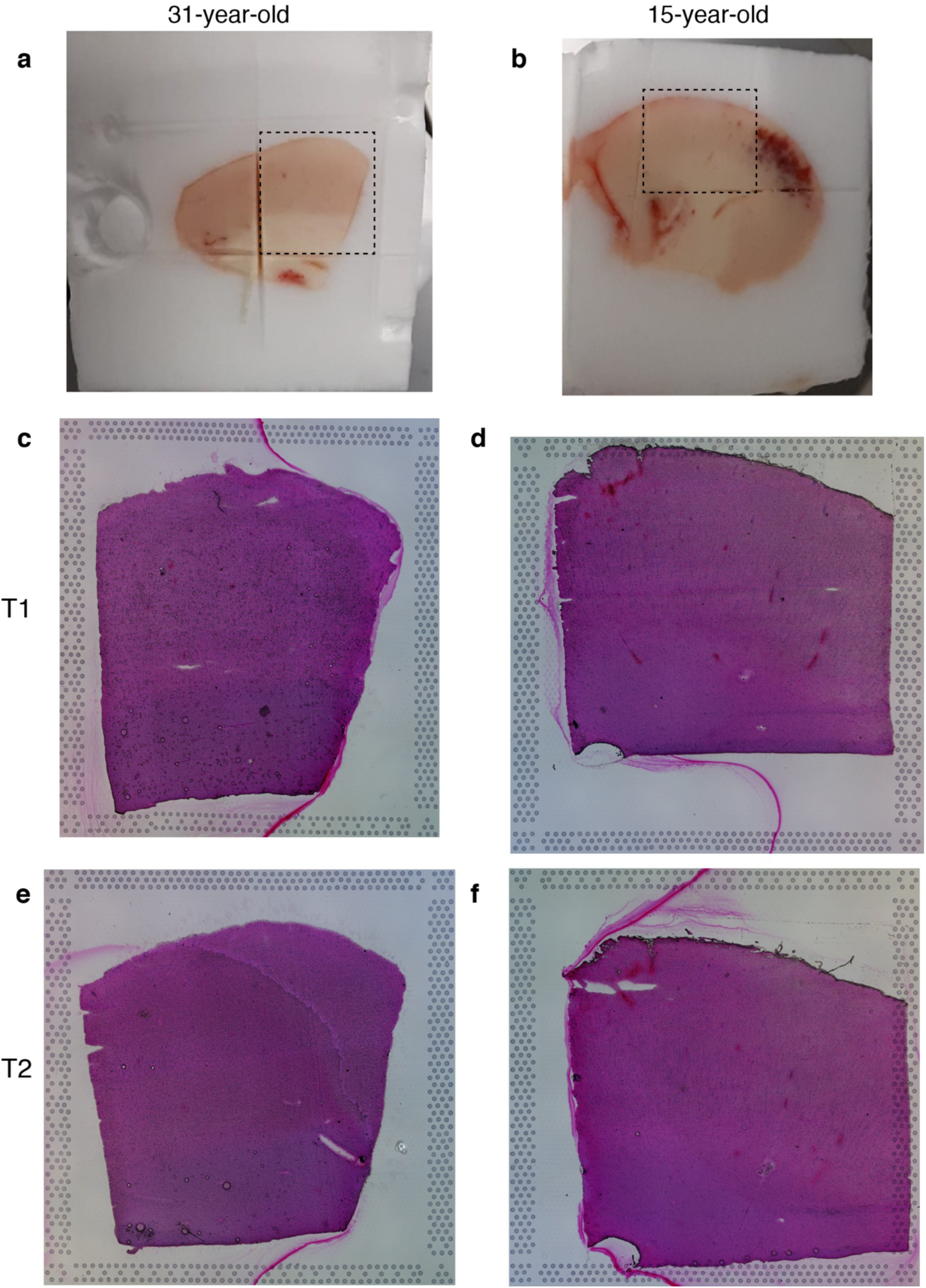
Visium Spatial Gene Expression samples. **a**,**b**, 31-year-old (**a**) and 15-year-old (**b**) temporal cortex tissue blocks embedded in OCT. Black dashed boxes outline the regions collected onto the Visium Spatial Gene Expression slide. **c**-**f**, H&E stained technical replicate tissue sections used to generate Visium Spatial Gene Expression libraries for the 31-year-old (**c,e**) and 15-year-old (**d,f**) tissue samples. T, technical replicate.

**Extended Data Fig. 5:**
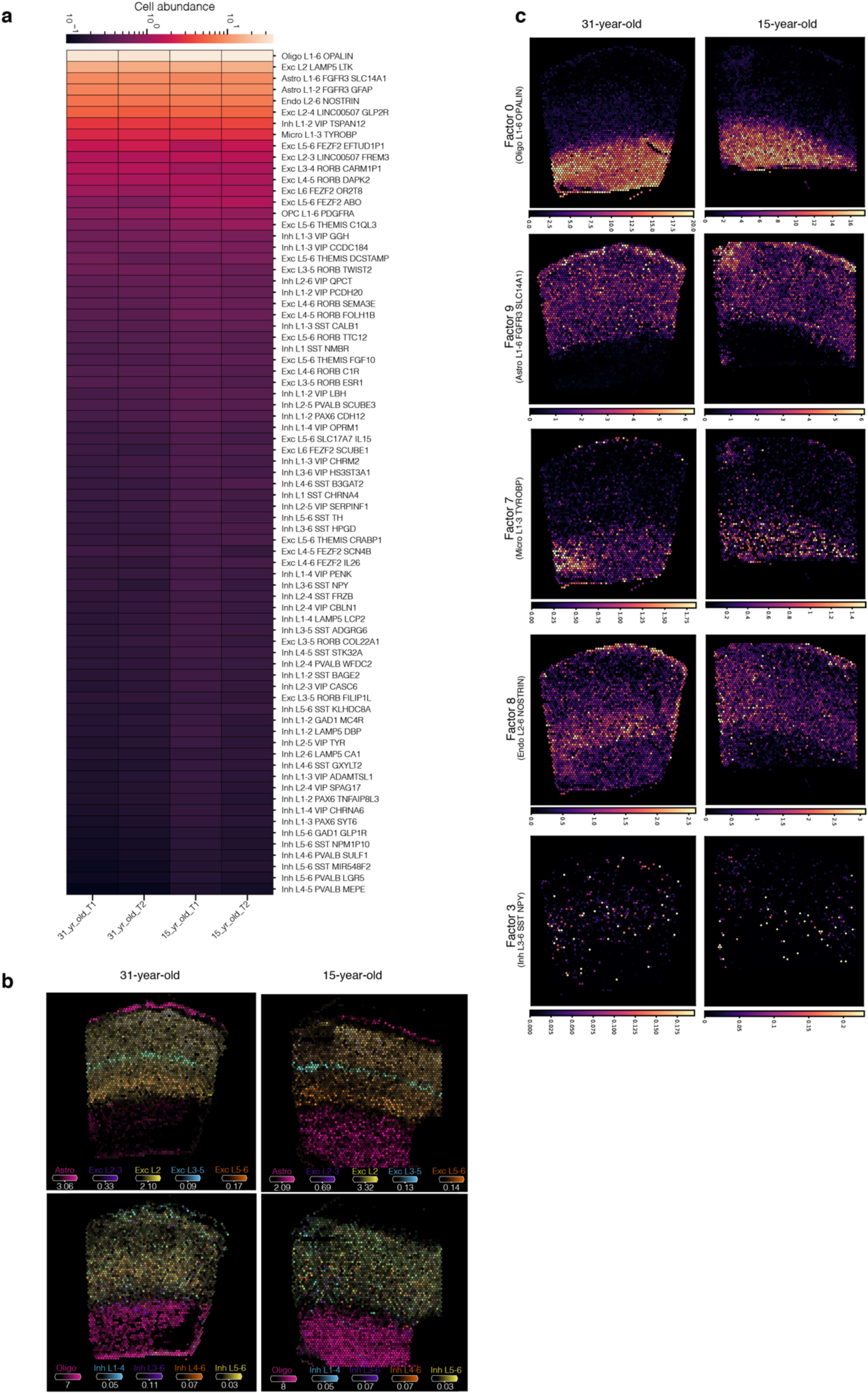
Spatial mapping of cell types in the human temporal cortex. **a,** Estimated cell abundance of 75 cell types across all Visium samples. Shown is a heatmap with the colour indicating the relative cell abundance of cell types (rows) across the different samples (columns). **b**, Estimated cell type abundances (colour intensity) in the technical replicate 31-year-old and 15-year-old temporal cortex tissue sections for a selection of cell types including non-neuronal cell types, excitatory neurons (top row) and inhibitory neurons (bottom row). **d**, Spatial plots show of the NMF weights for selected NMF factor/tissue compartment across the 31-year-old and 15-year-old temporal cortex tissue sections. Panels are displayed in the same order as the dotplot in Fig. 2c, with the dominant cell types for each factor indicated in brackets. T, technical replicate.

**Extended Data Fig. 6:**
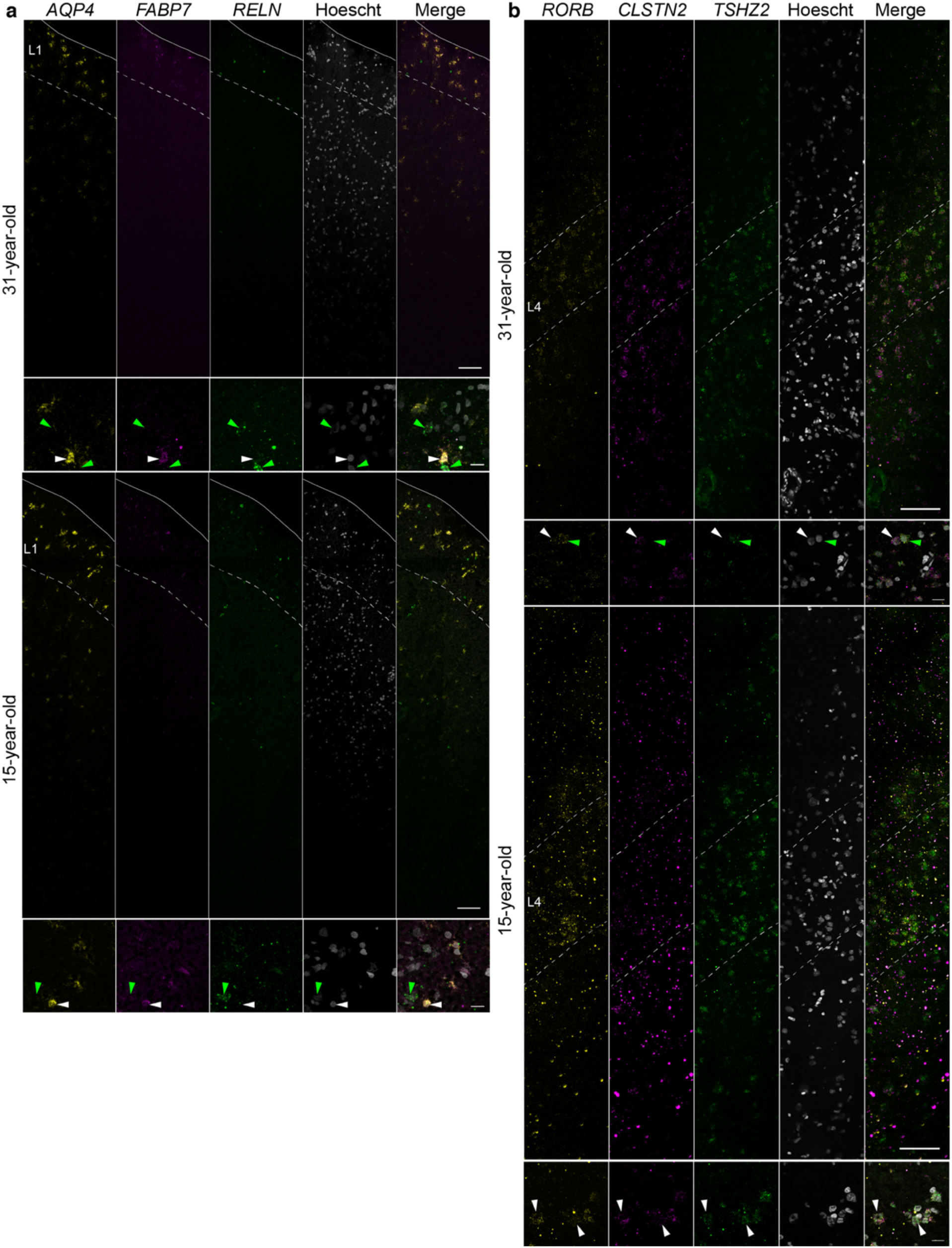
*In situ* HCR analysis of selected cortical layer marker genes. Expression of **a**, layer 1 markers *AQP4*, *FABP7* and *RELN* and **b**, layer 4-6 markers *RORB*, *CLSTN2* and *TSHZ2* in frozen temporal cortex tissue sections from the same 31-year-old and 15-year-old donor tissue used for Visium. High magnification views of layer 1 in **a** indicate *AQP4*/*RELN*-positive cells (yellow arrowheads) and *FABP7* positive cells (green arrowhead). In high magnification views of layer 4 in **b** in the 31-year-old tissue section, *RORB*/*CLSTN2*-positive (white arrowhead) and *RORB*/*TSHZ2*-positive cells (green arrowhead) are indicated. In high magnification views of layer 4 in **b** in the 15-year-old tissue section *RORB*/*CLSTN2*/*TSHZ2*-positive cells (white arrowheads) are indicated. Dashed white lines indicate layer boundaries. Solid white line indicates tissue edge. Scale bars are 100 µm in low magnification views (tile scan at 40x) and 20 µm in high magnification views (63x).

**Extended Data Fig. 7:**
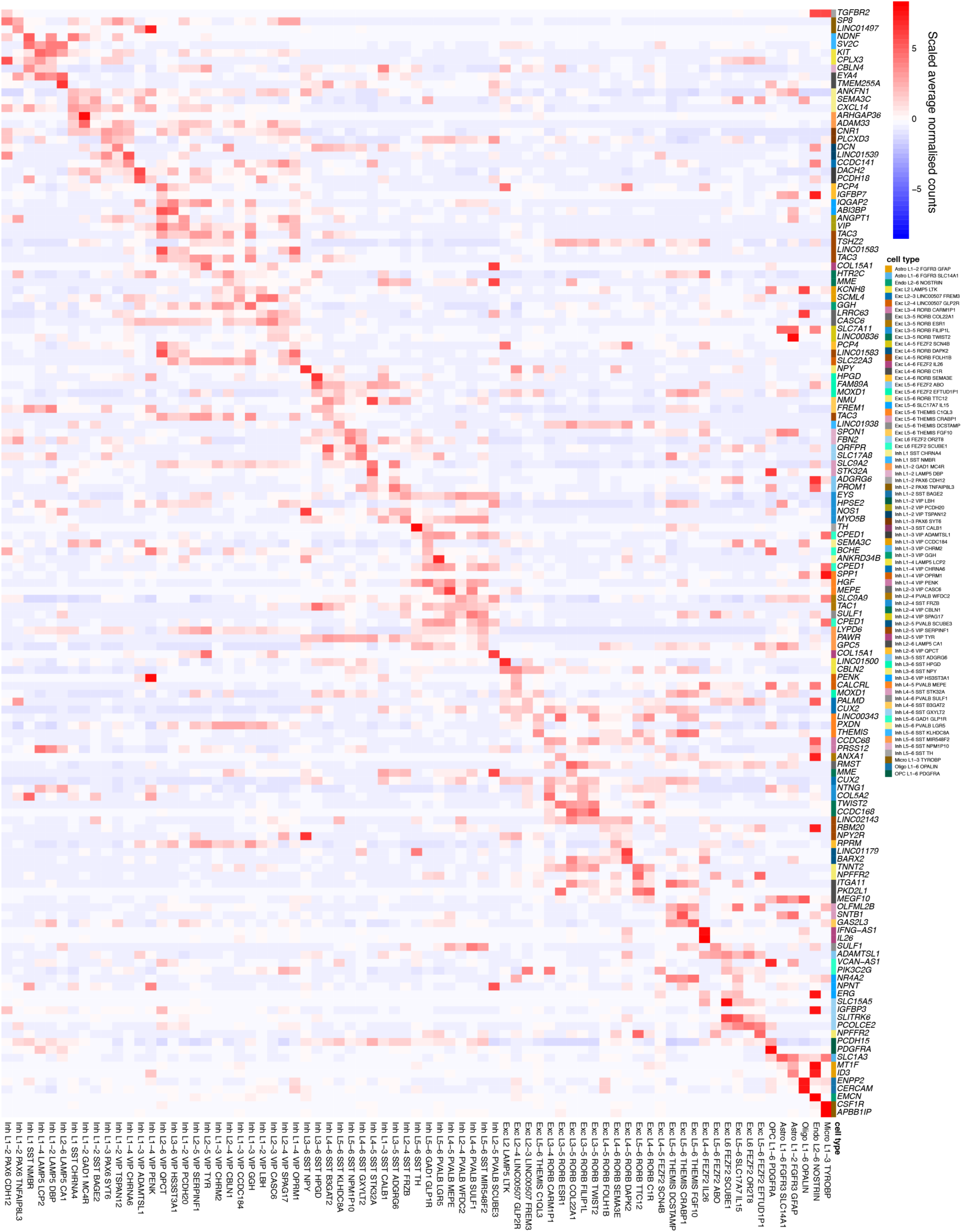
Expression of the reference MTG atlas minimal markers. Heatmap showing the scaled average normalised expression counts of the NS-Forest minimal marker genes identified for the reference MTG cell atlas dataset (y-axis) in each of the 75 query cortical cell types identified in the combined adult and paediatric snRNA-seq datasets (x-axis). The minimal marker genes are annotated (colour codes on the y-axes) according to the cell type they define.

**Extended Data Fig. 8:**
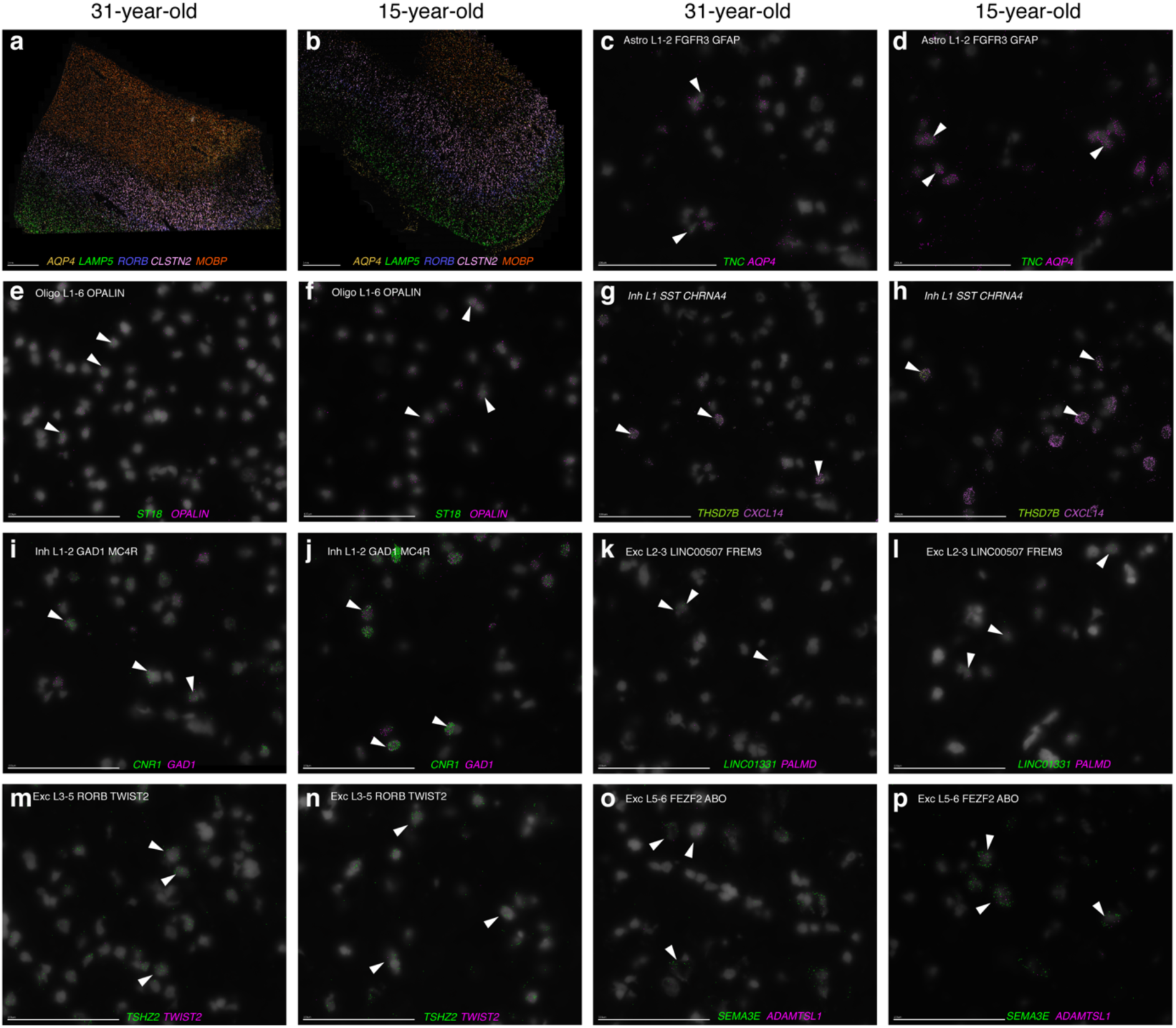
MERFISH spatial transcriptomics analysis of selected NS-forest markers. **a**, **b** Low magnification views of the 31-year-old (a) and 15-year-old (b) MERFISH datasets showing the expression of known layer maker genes in the expected layers as validation of the MERFISH experiment. **c**-**p**, High magnification views of 31-year-old (c,e,g,I,k,m,o) and 15-year-old (d,f,h,j,l,n,p) MERFISH datasets showing the overlap of new NS-Forest minimal markers (green) with published NS-Forest minimal markers (magenta) in indicated cells (arrowheads). The cell type that the NS-Forest markers are associated with is indicated in the top left corner. Scale bars: 100 µm.

**Extended Data Fig. 9:**
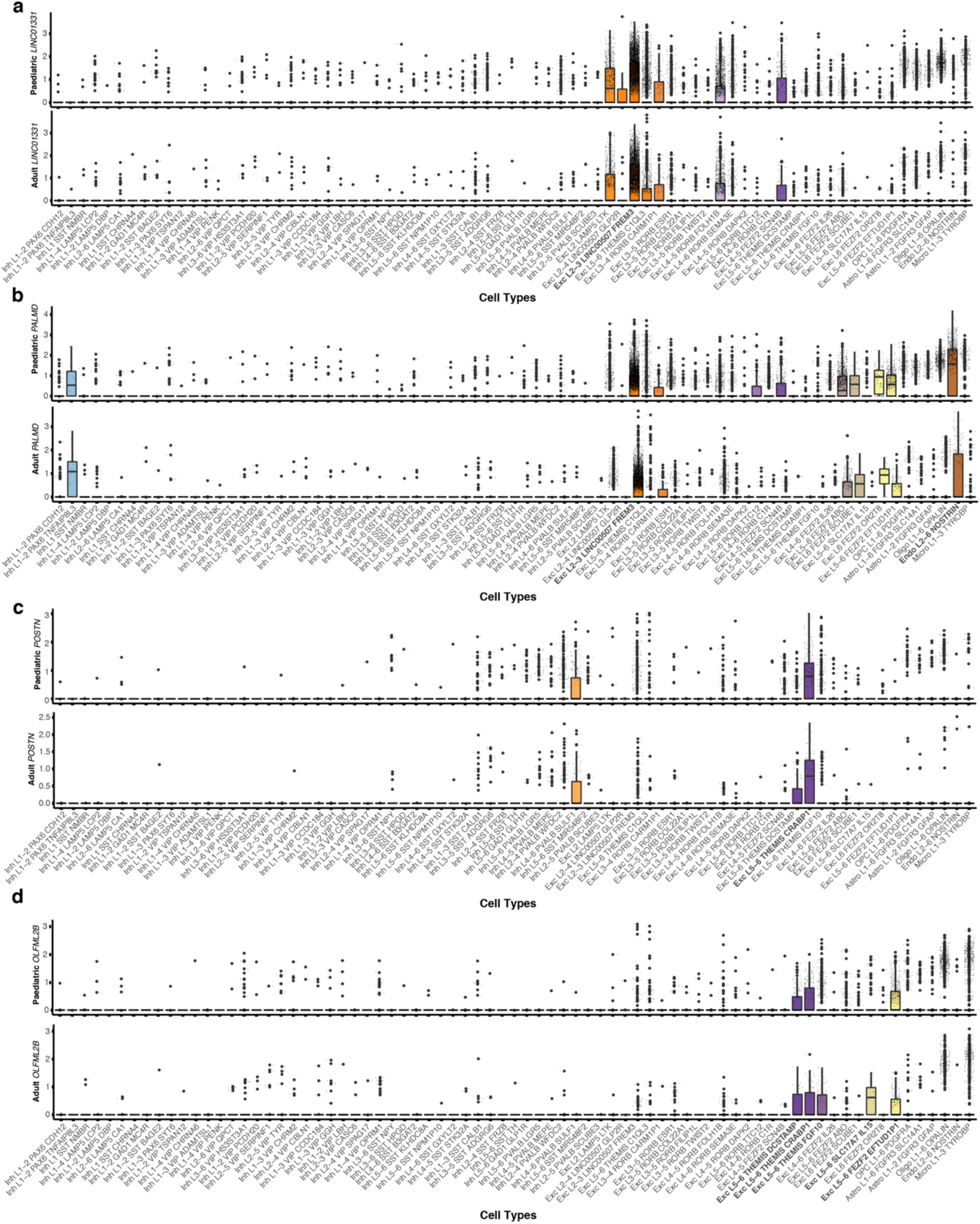
Evaluation of NS-Forest minimal marker gene expression across cell types in comparison to MTG cell taxonomy markers. **a-d**, Boxplots showing the normalised expression counts for *LINC01331* (**a**)*, PALMD* (**b**)*, POSTN* (**c**) and *OLFML2B* (**d**) in paediatric (top) and adult (bottom) datasets. The cell types expressing the markers at high levels are indicated in bold.

**Extended Data Fig. 10:**
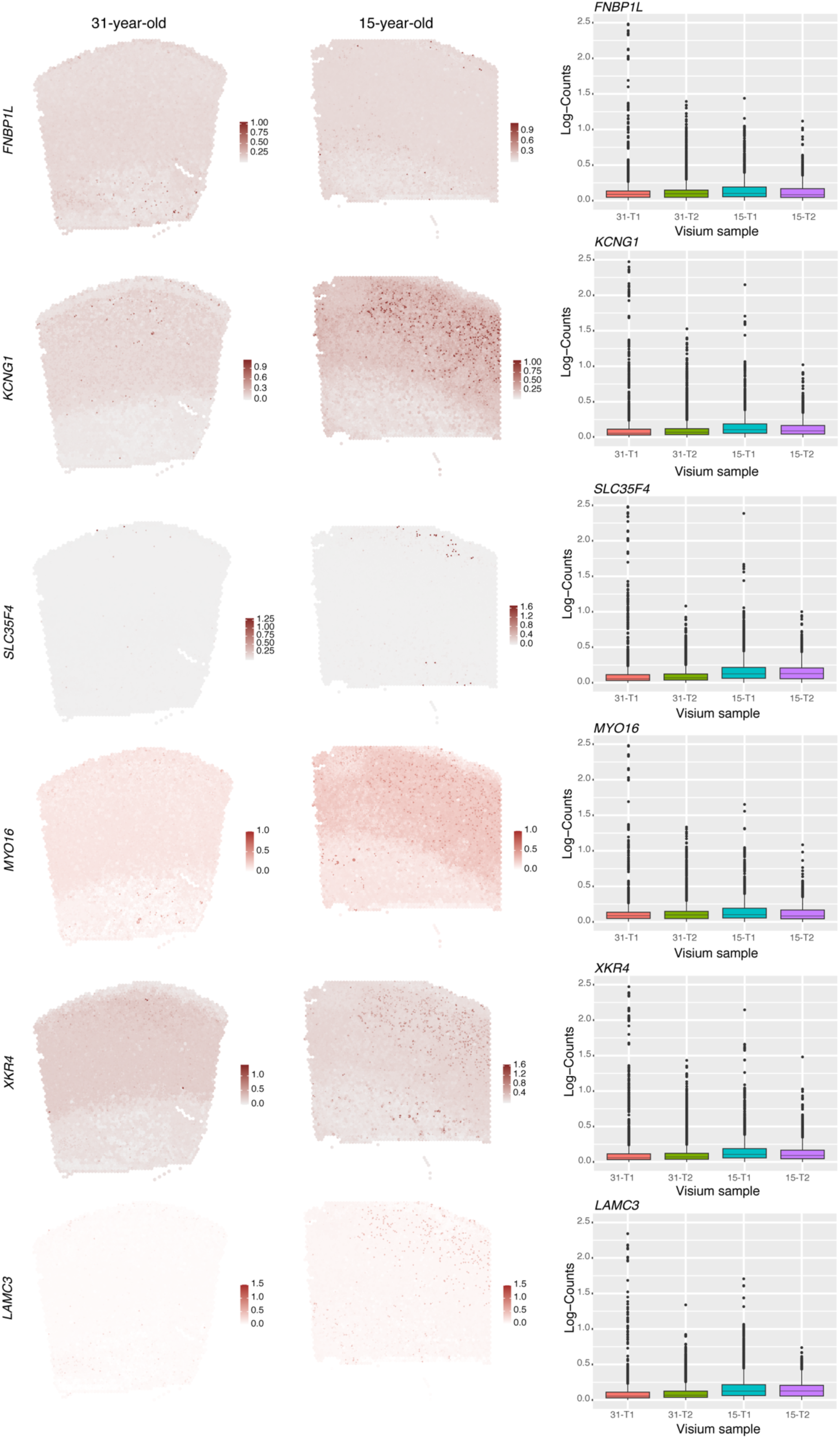
BayesSpace analysis of differentially expressed genes. High resolution Visium spatial gene expression profiles for selected DEGs using BayesSpace analysis to compare sub-spot level expression intensities between 31-year-old and 15-year-old temporal cortex tissue sections. Barplots show the average gene gene expression (log-counts) across technical replicate (T) samples for the indicated genes for spots with gene expression levels > 0. In all cases, average gene expression is higher in the 15-year-old samples than in the 31-year-old samples.

**Extended Data Fig. 11:**
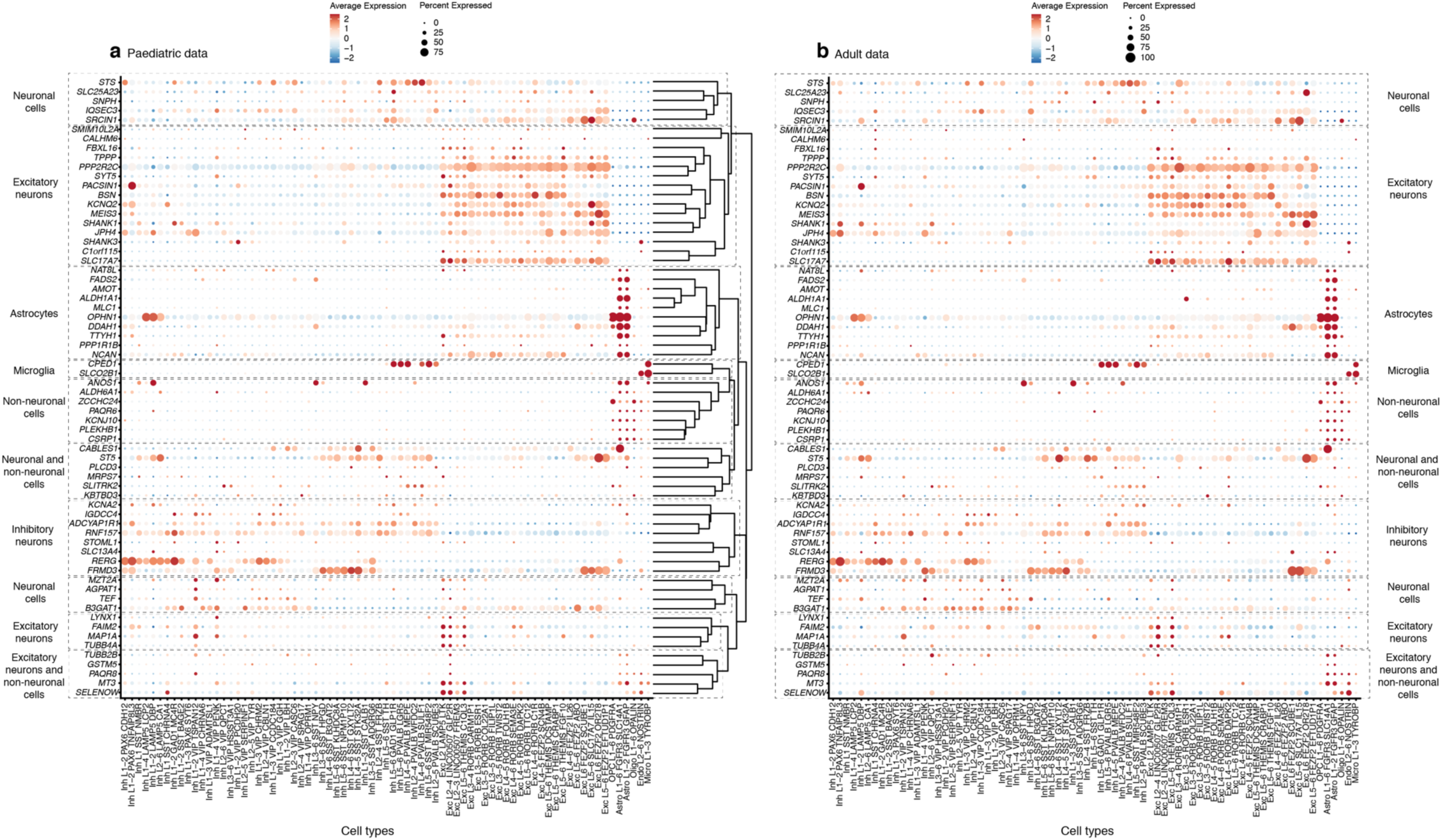
Cell type-specific expression of putative TBM biomarkers. **a**. Hierarchical clustering of TBM biomarker genes across the 75 cell types identified in the peaditaric snRNA-seq dataset reveals clusters of genes that are expressed by specific groups of cell types. **b**. Analysis of the same genes across the adult snRNA-seq dataset, using the gene order in (a) reveals very similar patterns of cell type-specific expression across the age-groups. Dashed boxes highlight gene clusters, with associated cell types indicated on the left and right of the right diagram

**Supplementary Fig. 1:**
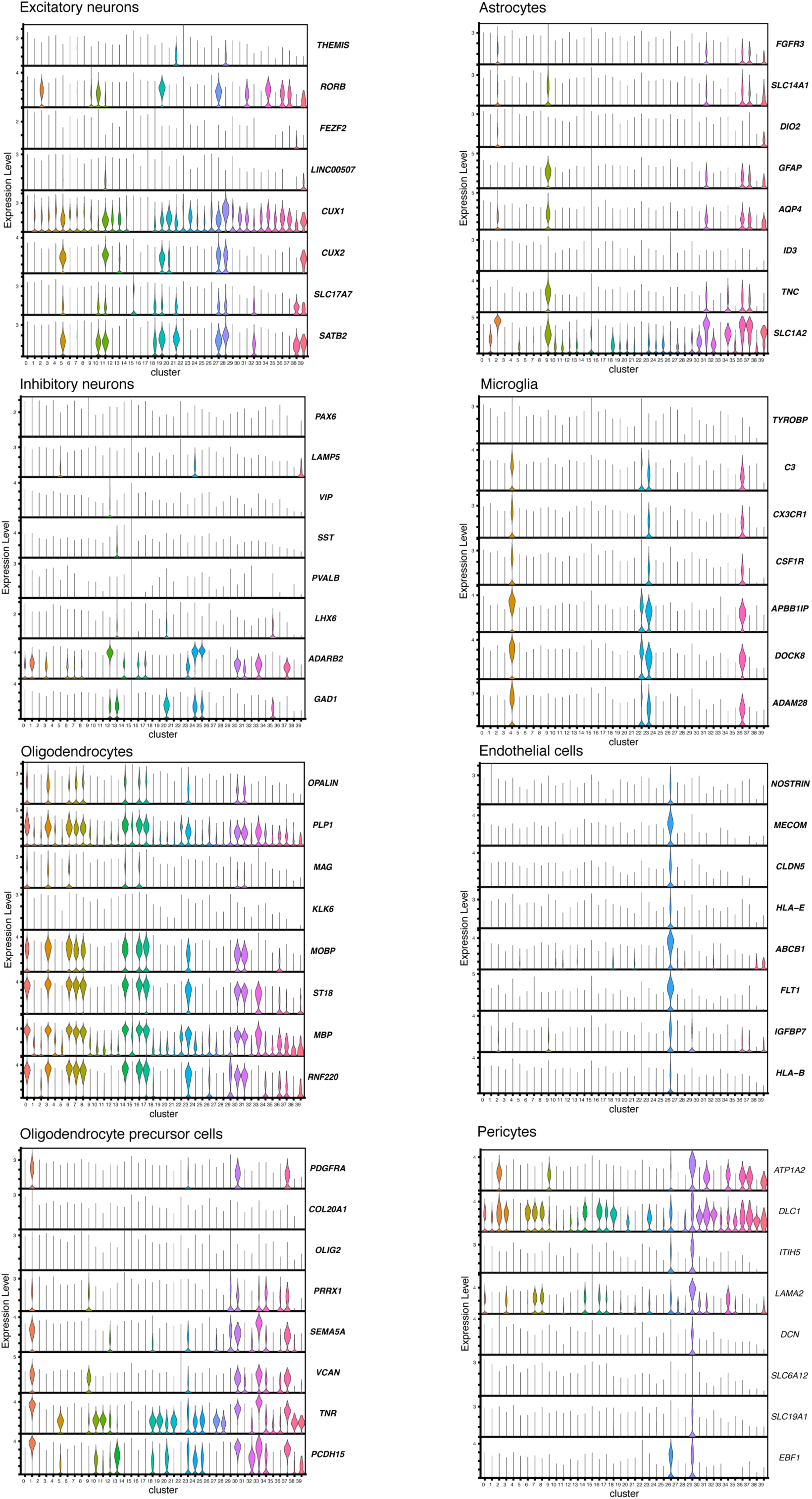
Violin plots showing the expression of known cell type marker genes across the Seurat clusters. These data were used for level 1 annotation of each cluster as one of the indicated major brain cell types (see also Extended Data Table 3).

**Supplementary Fig. 2:**
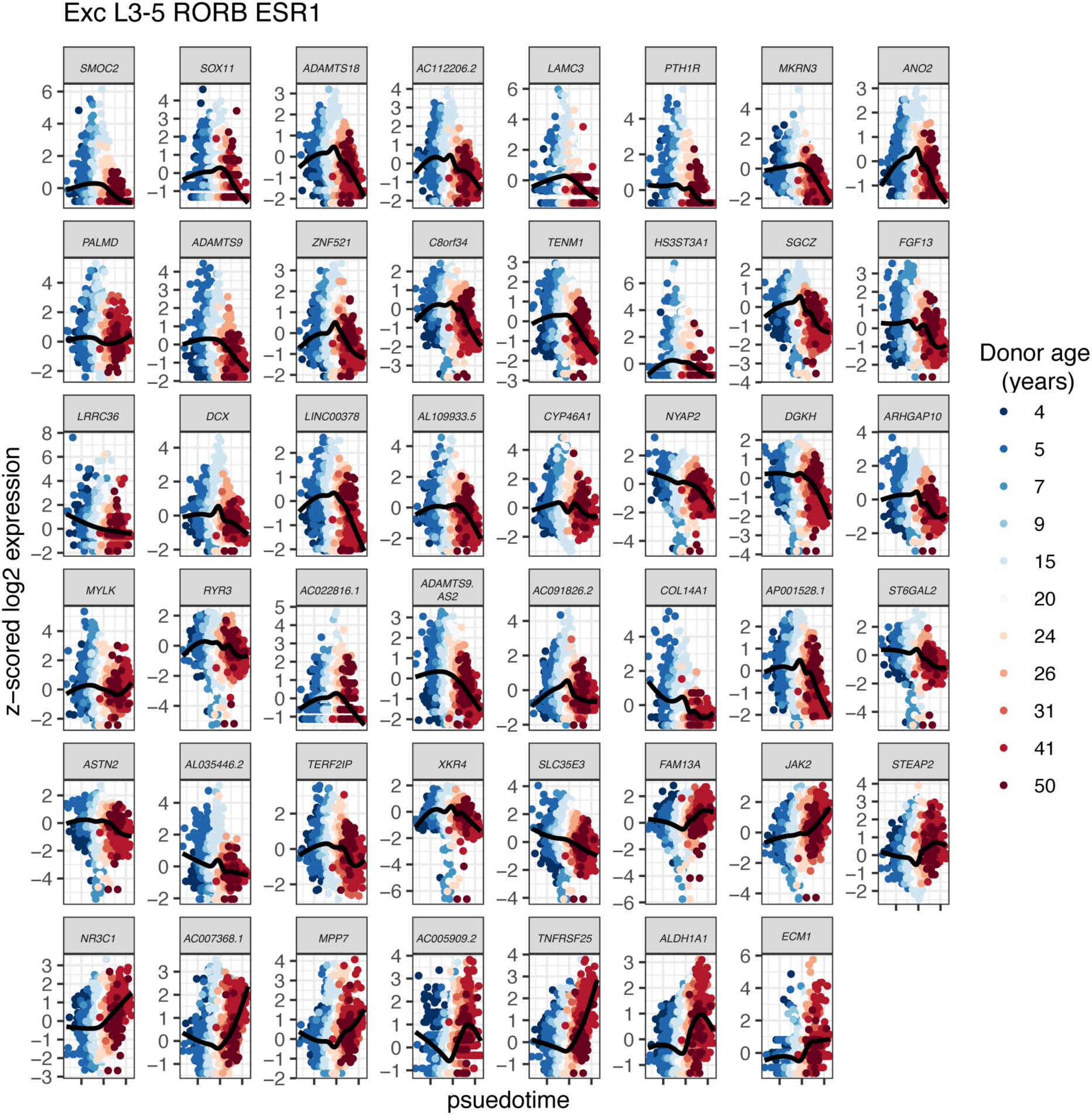
psupertime gene expression trajectories for all DEGs in Exc_L3-5_RORB_ESR1. The *x*-axis is the calculated psupertime value for each cell, coloured by sample of origin. The black lines are smoothened curves fit by geom_smooth in the R package ggplot2.

**Supplementary Fig. 3:**
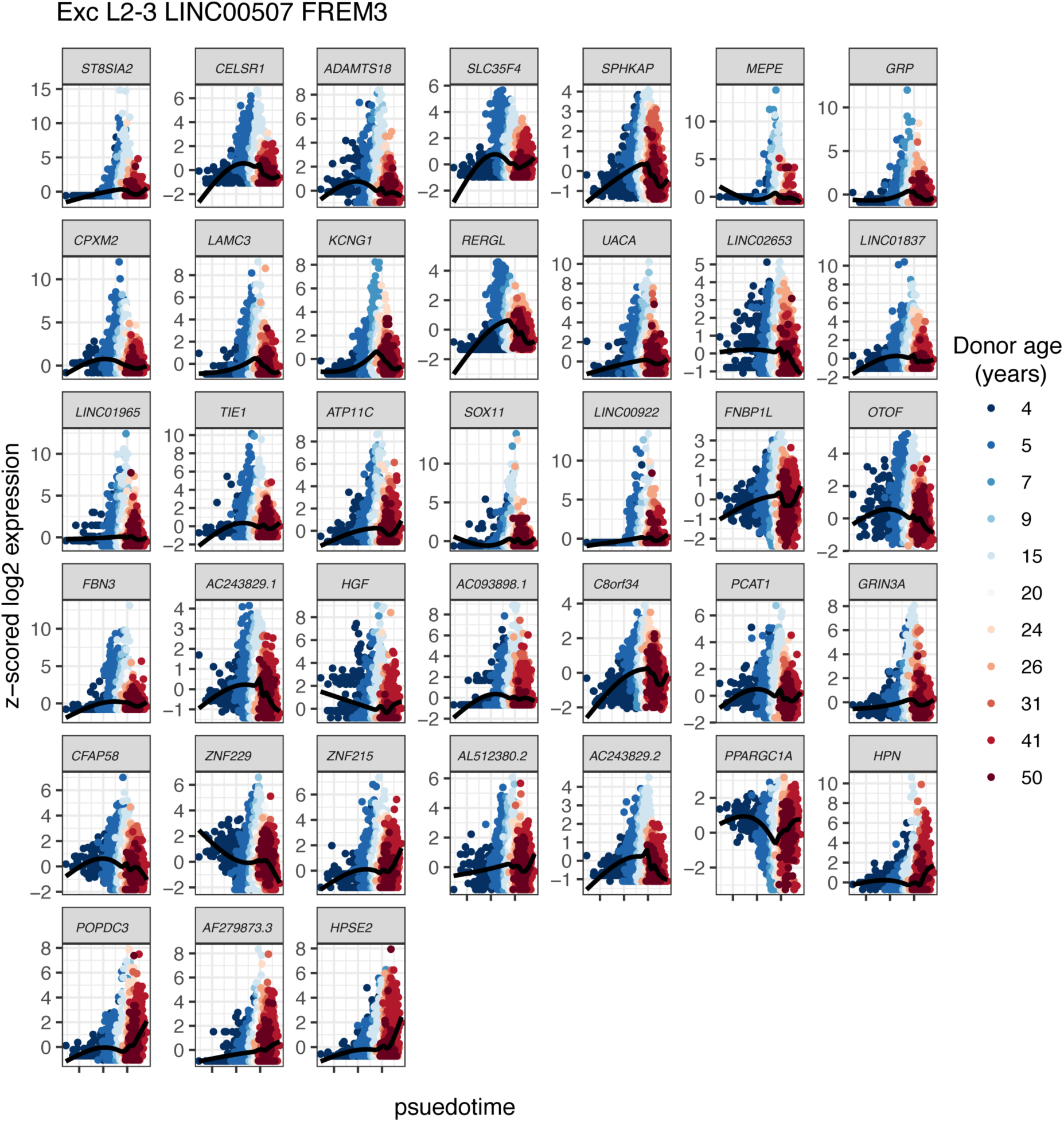
psupertime gene expression trajectories for all DEGs in Exc L2-3_LINC00507_FREM3. The *x*-axis is the calculated psupertime value for each cell, coloured by sample of origin. The black lines are smoothened curves fit by geom_smooth in the R package ggplot2.

**Supplementary Fig. 4:**
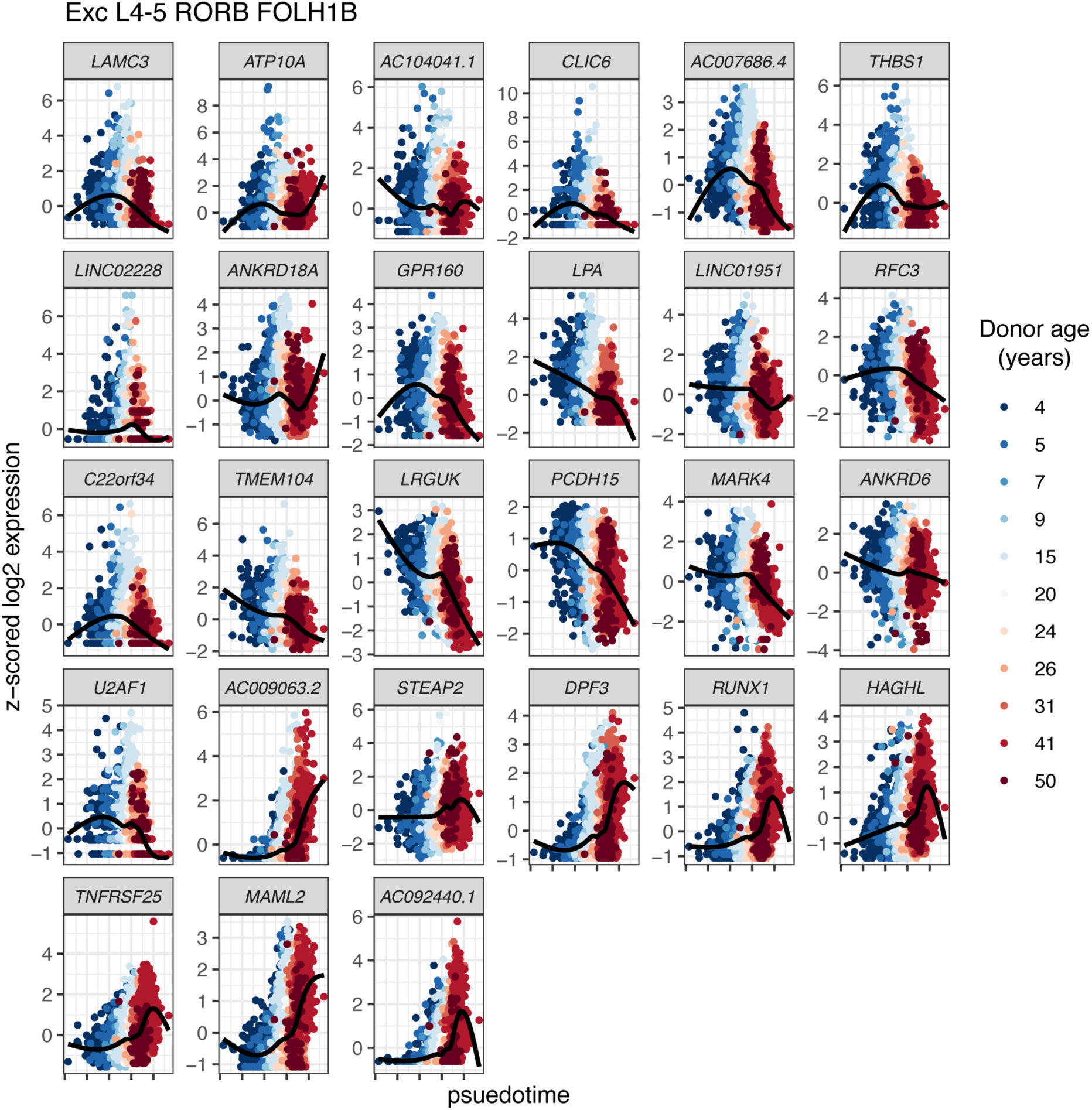
psupertime gene expression trajectories for all DEGs in Exc_L4-5_RORB_FOLH1B. The *x*-axis is the calculated psupertime value for each cell, coloured by sample of origin. The black lines are smoothened curves fit by geom_smooth in the R package ggplot2.

**Supplementary Fig. 5:**
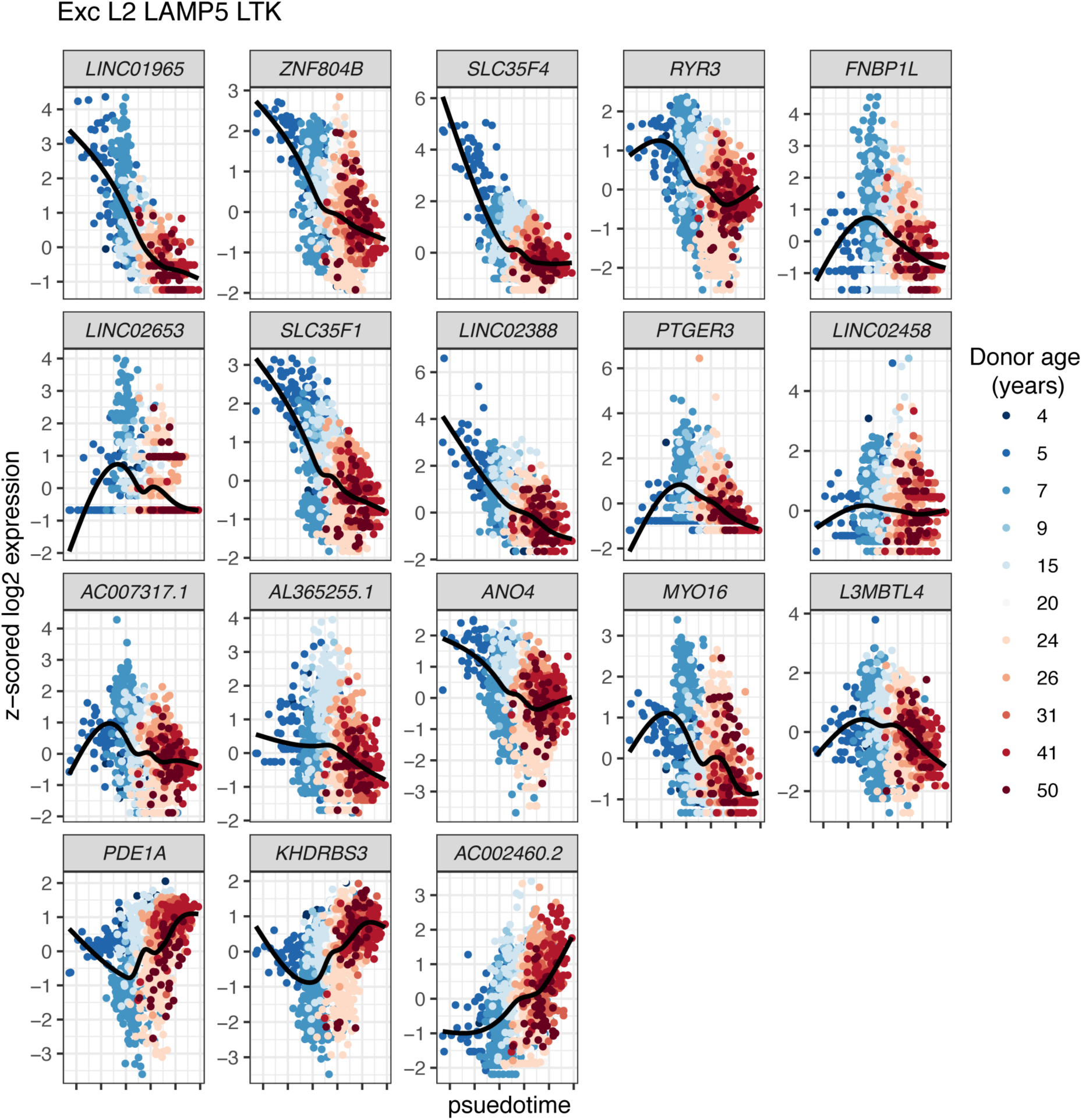
psupertime gene expression trajectories for all DEGs in Exc_L2_LAMP5_LTK. The *x*-axis is the calculated psupertime value for each cell, coloured by sample of origin. The black lines are smoothened curves fit by geom_smooth in the R package ggplot2.

## Extended data Tables

**Extended Data Table 1: Summary of snRNA-seq, Visium and MERFISH sample metadata.** Samples are ordered by age. The eight “P00” datasets were generated in the Hockman laboratory while the four “Nuc” datasets were generated by Thrupp et al. (2020)^24^.

**Extended Data Table 2: Summary of average quality control metrics for snRNA-seq datasets across nuclei for each sample before and after filtering.** Several measures for quality control were evaluated on a per sample basis including the sequencing saturation, the mean number of reads per nucleus, the number of barcodes, the median number of genes detected per nucleus, the median number of UMIs detected per nucleus, and the number of doublets removed.

**Extended Data Table 3: Label transfer annotation of snRNA-seq datasets using the Allen Brain Map MTG atlas as a reference. Sheet 1,** Manual annotation of clusters into major cell types (level 1 annotation). **Sheet 2**, Number of nuclei per level 1 annotation category per sample. **Sheet 3**, Number of nuclei per MTG cell type per sample. The number of barcodes corresponding to each MTG cell type and sample is shown. Additionally, the total, minimum, and maximum number of nuclei per cell type and sample was computed. The number of cell types represented per sample was also determined. **Sheet 4**, Number of removed nuclei per level 1 annotation category per sample. **Sheet 5**, Number of removed nuclei per MTG cell type per sample. **Sheet 6**, Subtraction matrix comparing cosine similarity scores (i.e. similarity score for each cell subtype compared to the MTG cell Atlas as in Fig. 1d) for paediatric dataset to the adult dataset. Values are the paediatric scores minus the adult scores. **Sheet 7**, p-values, tests performed for each cell type and padj values (Benjamini-Hochberg method) when comparing the proportion of nuclei between male and female samples. **Sheet 8-10,** p-values, tests performed for each cell type and padj values (Benjamini-Hochberg method) when comparing the proportion of nuclei (sheet 8), number of genes (sheet 9; see Extended Data Fig. 3b) and number of UMIs (sheet 10; see Extended Data Fig. 3c) for each cell type between paediatric and adult samples shown.

**Extended Data Table 4: Summary of average quality control metrics for Visium datasets.** Several measures for quality control were evaluated on a per sample basis including the sequencing saturation, the percentage of read mapped to the transcriptome, the number of spots under the tissue, the average number of nuclei per spot determined by Vistoseg analysis, the mean reads detected per spot, the median genes detected per spot, the total number of genes detected, the median UMI Counts per Spot and the total number of nuclei.

**Extended Data Table 5: NS-Forest minimal marker analysis. Sheet 1,** Statistical tests evaluating the expression of Aevermann et al. (2021)^29^ minimal markers (see their Supplementary Tables 1-2) in our datasets. **Sheets 2-3**, Metadata for each feature identified by NS-Forest marker in the down-sampled paediatric (**sheet 2**) and down-sampled adult (**sheet 3**) datasets describing the cell type, the F-beta score for each marker gene, overlap with Aevermann et al. (2021) and Hodge et al. (2019), uniqueness to the age group of interest, coding status, and uniqueness to the associated cell type as shown in Fig. 3. As input to NS-Forest, all datasets (six paediatric and six adult) were randomly down-sampled such that the total number of nuclei per sample was equal to the sample with the fewest number of nuclei.

**Extended Data Table 6: gProfiler analysis of NS-forest markers**. **Sheet1-3**, Significantly enriched GO terms associated with the paediatric (**sheet1**), adult (**sheet2**) and paediatric plus adult minimal marker genes identified by NS-forest. **Sheet3-5**, Significantly enriched GO terms associated with shared (i.e associated with both adult and paediatric samples) or paediatric-specific minimal marker genes with a binary expression score (> 0.7) for Oligo L1-6 OPALIN. Terms for which “highlighted” is true are driver terms.

**Extended Data Table 7: Summary of metadata for NS-Forest markers with a binary expression score (> 0.7) per cell type across the paediatric and adult datasets.** The number of shared markers, the number of markers unique to paediatric samples, and the number of markers unique to adult samples is shown for each cell type. The number of nuclei per cell type is shown for the combined paediatric and adult down-sampled datasets, the down-sampled paediatric datasets, and down-sampled adult datasets.

**Extended Data Table 8: Overlap NS-Forest markers with a binary expression score (> 0.7) per cell type between the paediatric and adult datasets.** NS-Forest markers with a binary expression score (> 0.7) per cell type were extracted for the down-sampled paediatric and down-sampled adult datasets. Each sheet represents 1 of 75 cortical cell types and the NS-Forest features which were shared (intersect) between the paediatric and adult datasets, unique to paediatric datasets, or unique to adult datasets are shown.

**Extended Data Table 9: DESeq2 output of all genes tested for differential expression between paediatric and adult brains per cell type. Sheet 1-75**, Differential expression analysis was performed using DESeq2’s Wald Test for each cell type separately. Genes were filtered prior to testing to only include those expressed in > 10% of nuclei for that cell type across all paediatric and adult datasets. The associated log2FoldChanges, p-adjusted values (padj, Benjamini-Hochberg method), and description of each feature are shown. Positive log_2_FoldChanges represent genes upregulated in paediatrics versus adults. See DESeq2 documentation for explanation of NA values (https://bioconductor.org/packages/release/bioc/vignettes/DESeq2/inst/doc/DESeq2.html#why-are-some-p-values-set-to-na).

**Extended Data Table 10: DESeq2 output of significant DEGs only between paediatric and adult brains in a subset of cell types. Sheet 1-21**, Significant DEGs (padj < 0.05) for cell types shown in Fig. 5a. The associated log_2_FoldChanges, p-adjusted values (padj), description, percentage of paediatric nuclei expressing the gene, percentage of adult nuclei expressing the gene, average normalised expression across paediatric nuclei, and average normalised expression across adult nuclei are shown. The difference in the peaditaric and adult values for percentage of nuclei and average normalised expression is also shown. Positive log_2_FoldChanges represent genes upregulated in paediatric versus adults datasets. See DESeq2 documentation for explanation of NA values (https://bioconductor.org/packages/release/bioc/vignettes/DESeq2/inst/doc/DESeq2.html#why-are-some-p-values-set-to-na). **Sheet 22-24**, EA (sheet 6), IQ (sheet7) and HAR (sheet8) associated DEGs and their associated cell types.

**Extended Data Table 11: psupertime coefficients.** The calculated psupertime coefficients for each gene for indicated excitatory neuron subtypes that showed the highest number of DEGs. Genes with non-zero psupertime coefficients represent genes that are relevant to the ordering of the cells in pseudotime.

**Extended Data Table 12**: **GSEA terms associated with each cell type showing enriched or depleted pathways in paediatric versus adult samples.** GSEA was performed using DESeq2’s output gene lists for each cell type ranked according to the log2FoldChange*-log_2_(padj) for each gene. All DESeq2-tested genes served as input into GSEA (genes were expressed in > 10% of nuclei for the cell type of interest). Matrix shows the corresponding positive (**sheet 1**) and negative (**sheet 2**) NES values for each GSEA term (y axis) and cell type (x axis) based on the analysis using the ranked list of genes for each cell type. Terms were filtered to only include significantly associated terms (p<0.01, q<0.1). Positive NES values indicate pathways that are enriched in paediatric versus adult samples; negative NES values indicate pathways that are depleted in paediatric versus adult samples. The total number of terms per cell type and the total number of cell types associated with a given term are shown.

